# Developing an anatomically valid segmentation protocol for anterior regions of the medial temporal lobe for neurodegenerative diseases

**DOI:** 10.1101/2025.02.11.637506

**Authors:** Niyousha Sadeghpour, Sydney A. Lim, Anika Wuestefeld, Amanda E. Denning, Ranjit Ittyerah, Winifred Trotman, Eunice Chung, Shokufeh Sadaghiani, Karthik Prabhakaran, Madigan L. Bedard, Daniel T. Ohm, Emilio Artacho-Pérula, Maria Mercedes Iñiguez de Onzoño Martin, Monica Muñoz, Francisco Javier Molina Romero, José Carlos Delgado González, María del Mar Arroyo Jiménez, Maria del Pilar Marcos Rabal, Ana María Insausti Serrano, Noemí Vilaseca González, Sandra Cebada Sánchez, Carlos de la Rosa Prieto, Ricardo Insausti, Corey McMillan, Edward B. Lee, John A. Detre, Sandhitsu R. Das, Long Xie, M. Dylan Tisdall, David J. Irwin, David A. Wolk, Paul A. Yushkevich, Laura EM. Wisse

**Affiliations:** Department of Radiology, University of Pennsylvania, Philadelphia, Pennsylvania, USA; Department of Clinical Sciences, Lund University, Malmö, Sweden; Department of Neurology, University of Pennsylvania, Philadelphia, Pennsylvania, USA; Department of Pathology and Laboratory Medicine, University of Pennsylvania, Philadelphia, Pennsylvania, USA; Human Neuroanatomy Laboratory, Neuromax CSIC Associated Unit, University of Castilla-La Mancha and Institute for Biomedicine, Albacete, Spain; Department of Digital Technology and Innovation, Siemens Healthineers, Princeton, NJ 08540, USA; Department of Clinical Sciences, Lund University, Lund, Sweden

**Keywords:** Medial temporal lobe, Alzheimer’s disease, imaging biomarkers

## Abstract

**Background:** The anterior portion of the medial temporal lobe (MTL) is one of the first regions targeted by pathology in sporadic Alzheimer’s disease (AD) and Limbic-predominant Age-related TDP-43 Encephalopathy (LATE) indicating a potential for metrics from this region to serve as imaging biomarkers. Leveraging a unique post-mortem dataset of histology and magnetic resonance imaging (MRI) scans we aimed to 1) develop an anatomically valid segmentation protocol for anterior entorhinal cortex (ERC), Brodmann Area (BA) 35, and BA36 for in vivo 3 tesla (T) MRI and 2) incorporate this protocol in an automated approach.

**Methods:** We included 20 cases (61-97 years old, 50% females) with and without neurodegenerative diseases (11 vs. 9 cases) to ensure generalizability of the developed protocol. Digitized MTL Nissl-stained coronal histology sections from these cases were annotated and registered to same-subject post-mortem MRI. The protocol was developed by determining the location of histological borders of the MTL cortices in relation to anatomical landmarks. Subsequently the protocol was applied to 15 cases twice, with a 2-week interval, to assess intra-rater reliability with the Dice Similarity Index (DSI). Thereafter it was implemented in our in-house Automatic Segmentation of Hippocampal Subfields (ASHS)-T1 approach and evaluated with DSIs.

**Results:** The anterior histological border distances of ERC, BA35 and BA36 were evaluated with respect to various anatomical landmarks and the distance relative to the beginning of the hippocampus was chosen. To formulate segmentation rules, we examined the histological sections for the location of borders in relationship to anatomical landmarks in the coronal sections. The DSI for the anterior MTL cortices for the intra-rater reliability was 0.85-0.88 and for the ASHS-T1 against the manual segmentation was 0.62-0.65.

**Discussion:** We developed a reliable segmentation protocol and incorporated it in an automated approach. Given the vulnerability of the anterior MTL cortices to tau deposition in AD and LATE, the updated approach is expected to improve imaging biomarkers for these diseases.

## 1. Introduction

The medial temporal lobe (MTL) is a multifaceted and complex brain structure involved in memory function (2). It is characterized by its heterogeneity and comprises several cytoarchitectonically, connectomically and functionally distinct subregions (3). MTL subregions, including the entorhinal cortex (ERC), perirhinal cortex (PRC) and hippocampus, are differentially involved in different memory processes, such as recall and recognition (4–6), and are therefore highly relevant for memory decline in the context of neurodegenerative diseases. Not surprisingly, the MTL is a vulnerable region of pathology in various neurodegenerative diseases, including sporadic Alzheimer’s disease (AD) and Limbic-predominant Age-related TDP-43 Encephalopathy (LATE).These diseases are known to be associated with prominent deposition of pathology within this region (7,8) leading to neurodegeneration and cognitive decline. Notably, in the early phases of both AD and LATE, the involvement of the MTL is not uniform.

Rather, it exhibits changes in selective subregions, suggesting a potential for magnetic resonance imaging (MRI)-based morphometry in this region to be used as a valuable imaging biomarker.

Evidence suggests that in AD, tau deposition initially targets anterior portions of the MTL cortex, prominently affecting the PRC, particularly Brodmann area 35 (BA35)(7–9), also referred to as transentorhinal cortex(10), as well as the ERC. With disease progression, the pathology accumulates in the medial ERC and involves the cornu ammonis 1 (CA1) subfield of the hippocampus (8). In LATE, the amygdala is first targeted by TDP-43 pathology, followed by the ERC and hippocampus (7).

MTL imaging biomarkers, and especially hippocampal volume measured on MRI, have proven valuable as a diagnostic tool in memory clinics (11) and as an endpoint in clinical trials (12). However, with the increasing focus on identifying and treating early stages of dementia (13), there is a need for more precise MRI measures that can detect earliest changes, beyond total hippocampal volume, especially since the hippocampus is not the first affected region in either AD or LATE (7,8). MTL cortical measures, such as ERC and BA35 and especially metrics from the anterior regions, could be valuable biomarkers for this purpose.

However, these regions are challenging to measure because of the anatomical variability they exhibit (14). Manual segmentation protocols that include these MTL cortices and even the anterior portions are available (15,16). Nonetheless manual segmentation is labor intensive, time consuming and requires specialized expertise- factors that are prohibitive when dealing with larger datasets. Hence automated segmentation methods were introduced. Several automated segmentation methods exist that include MTL cortices (1,17–20). However, these methods either do not include the most anterior portions, lack fine-grained labels such as BA35, or fail to account for the anatomical variability of this region, either in the segmentation protocol or because of the use of a single-atlas approach. One of the automated approaches is the Automatic Segmentation of Hippocampal Subfields (ASHS) pipeline, which was first generated using high-resolution oblique coronal T2-weighted MRI images of the MTL (17). This pipeline includes fine-grained labels in the hippocampus and MTL cortex and, owing to the design of the protocol and the multi-atlas segmentation approach, accounts for anatomical variability in the MTL. More recently, Xie et al. (1) extended ASHS to more widely available 1x1x1mm^3^ resolution T1-weighted MRI images. This ASHS-T1 approach labels MTL cortical subregions similarly to the original ASHS, but does not label distinct hippocampal subfields, which are hard to distinguish in T1- weighted MRI at this resolution(21). Neither ASHS nor ASHS-T1 include the anterior portion of MTL which can be particularly vulnerable to tau tangles and TDP-43 pathology (7,22).

To address the limitations of available MTL segmentation tools and protocols with respect to the anterior MTL, our first objective was to develop an anatomically grounded and detailed manual segmentation protocol for the anterior ERC, BA35, and BA36 to be applied to *in vivo* T1-weighted 3 tesla (T) MRI. The rules for this protocol are based on the analysis of the location of subregion boundaries in a unique dataset of ultra-high-resolution post-mortem MRI and matched serial Nissl histology from 20 cases, annotated by expert neuroanatomists. Our second objective was to employ this new protocol to expand the existing ASHS-T1 atlas into the anterior MTL after assessing the intra-rater reliability of a manual rater. We then evaluated the cross-validation accuracy of ASHS-T1 for these newly added anterior MTL subregions.

## 2. Materials and Methods

See Fig. 1 for a flowchart of the different parts of protocol development and validation and the data used for each part.

**Figure 1.**
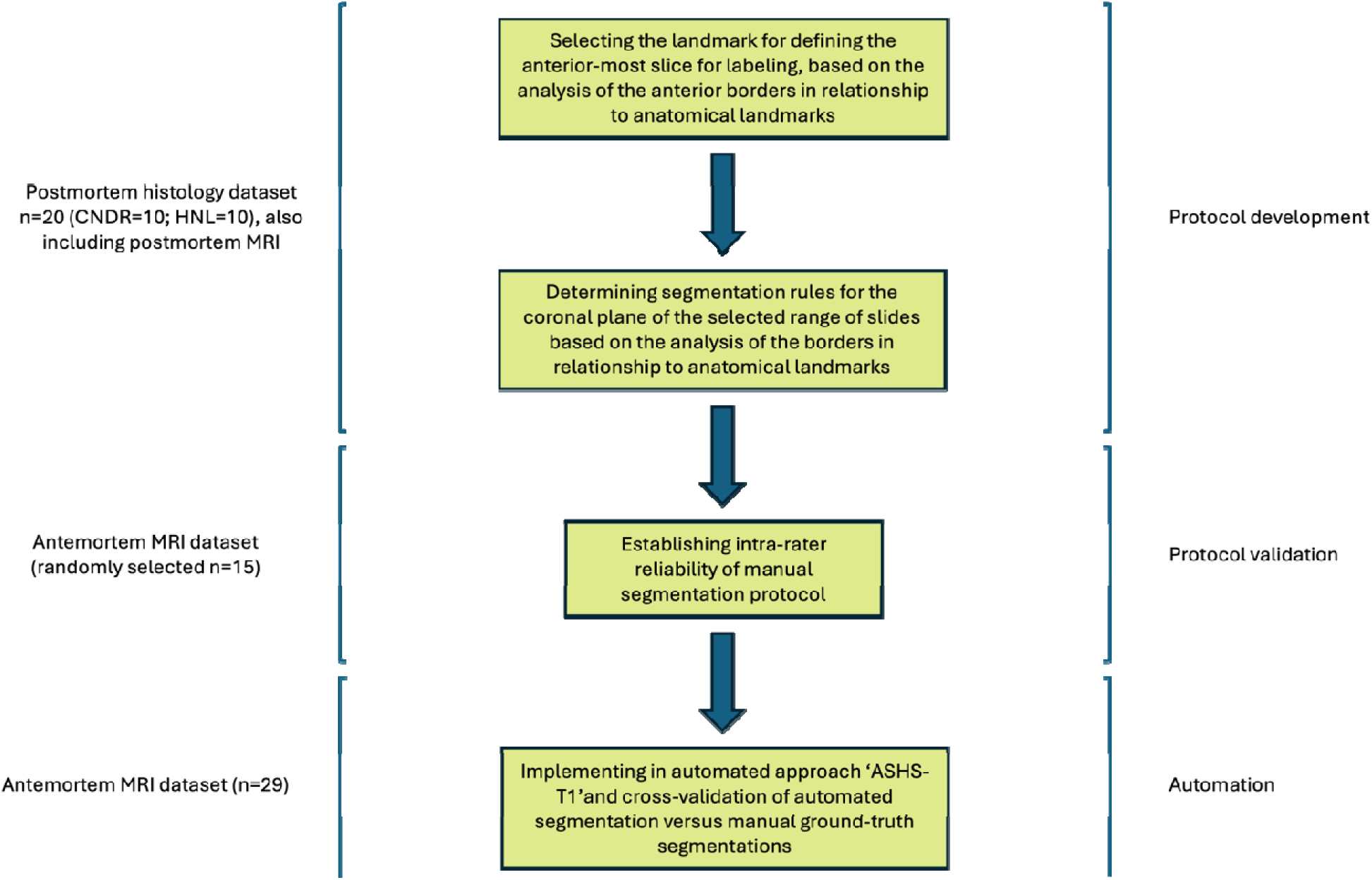
Flowchart of methods and datasets used in protocol development. This flowchart provides an overview of the key steps and datasets utilized in developing the segmentation protocol for anterior BA35, BA36 and ERC. Abbreviations: BA=Brodmann Area; ERC=entorhinal cortex; HNL: the Human Neuroanatomy Lab; CNDR: the Center for Neurodegenerative Disease Research.

### 2.1. *Ex vivo* population

We obtained brain hemisphere specimens from autopsy cases at the Human Neuroanatomy Lab (HNL) at the University of Castilla-La Mancha (UCLM), Spain, and the Center for Neurodegenerative Disease Research (CNDR) at the University of Pennsylvania, USA. Human brain specimens were obtained in accordance with local laws and regulations at the University of Pennsylvania and the Ethical Committee of UCLM. When possible, pre-consent was secured during the subjects’ lifetime. In all cases, consent from the next-of-kin was obtained after the subjects’ passing. Donors from CNDR included patients from the Penn Frontotemporal Degeneration Center and the Penn Alzheimer’s Disease Research Center participating in *in vivo* aging and dementia research. UCLM donors were mostly older adults with no known neurological diseases from the surrounding geographical area (23,24).

We selected 20 cases for the development of the manual segmentation protocol. To ensure generalizability of the protocol within older populations, the chosen autopsy cases were with and without neurodegenerative diseases (11 vs. 9 cases, respectively), with a relatively broad age range and an equal representation of men and women. Table 1 provides a summary of the demographic and diagnostic information for the cases we included, with additional details in Supplementary Table 1.

**Table 1.**
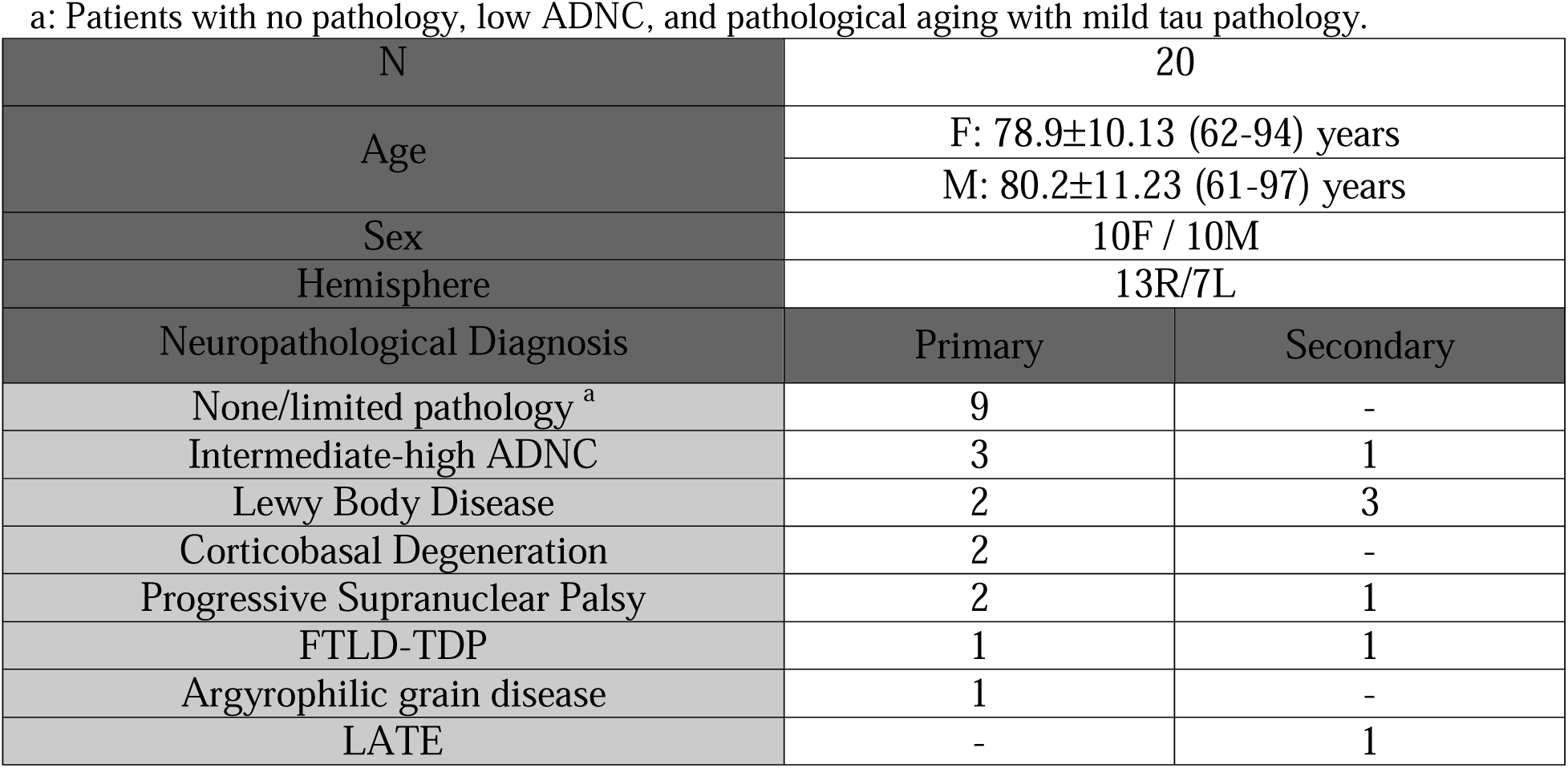
Demographic and diagnostic summary of the brain donors included in this study. Abbreviations: ADNC: Alzheimer’s disease neuropathologic change; LATE: Limbic Age-related TDP-43 Encephalopathy; FTLD-TDP: Frontotemporal Lobar Degeneration with TDP-43 inclusions a: Patients with no pathology, low ADNC, and pathological aging with mild tau pathology.

### 2.2. Histology data, annotation, and post-mortem MRI

#### 2.2 *Ex vivo* imaging procedure

For each of the brains donated, one hemisphere was used for imaging and serial histology preparation, while the opposite hemisphere was sampled for diagnostic pathology according to the National Institute on Aging–Alzheimer’s Association protocol (25,26). CNDR hemispheres were fixed in 10% formalin solution for at least 30 days. HNL hemispheres were fixed in situ with 8 L of 4% paraformaldehyde perfused through both carotid arteries, and then stored submerged in 4% paraformaldehyde for 6 weeks before being processed (27).

Following fixation, the temporal lobe was extracted from every hemisphere. The excised specimen was imaged overnight on a Varian 9.4 T animal scanner at 0.20 x 0.20 x 0.20 mm^3^ resolution using a multi-slice spin echo sequence. Sequence parameters vary slightly between specimens, with typical values being a repetition time of 9330 ms, and an echo time of 23 ms. Further details of the MRI acquisition and processing protocol can be found in Yushkevich et al. (23).

After MRI scanning, serial histological processing was performed on each MTL specimen (27). First, custom molds were 3-D printed to fit each temporal lobe specimen. The specimens were then cut into 2 cm blocks perpendicular to the long axis of the hippocampus using the molds. Next, the blocks were cryoprotected and sectioned coronally at 50 μm intervals using a sliding microtome coupled to a freezing unit (Microm, Heidelberg). As a result, each block generated ∼40 sections with every 10^th^ section undergoing staining for Nissl using a 0.25% Thionin stain.

Stained sections were then mounted on 75 mm x 50 mm glass slides, digitally scanned at pixel size 0.4µm at 20X magnification and uploaded to an open-source cloud-based digital histology archive (https://github.com/pyushkevich/histoannot) that supports web-based visualization, anatomical labeling, and machine learning classifier training. The MTL cortex was annotated in the histology sections by expert neuroanatomists, led by RI. For more details on the annotation protocol, please see the Supplementary Methods section.

Additionally, Supplementary Fig. 5 shows 3D renderings for two cases where the histological annotations were mapped into postmortem MRI. These 3D renderings show consistency in the borders of these anterior MTL cortical regions between consecutive slices.

### 2.3. Approach to protocol development

Since the goal of this work is to extend the existing segmentation protocol used in ASHS-T1 to more anterior cortical MTL regions, the MRI plane in which the tracing of these regions is performed, needed to match that of the existing protocol. In the ASHS-T1 protocol, this plane is approximately perpendicular to the main axis of the hippocampus. For brevity, we will refer to this plane as the “coronal” plane. The plane of histological sectioning was also chosen to be approximately perpendicular to the hippocampus main axis. By maintaining the same anatomical orientation for the MRI and serial histology we were able to use distances in the coordinate system of serial histology slices to inform rules for placing landmarks in the extended ASHS-T1 protocol.

An examination of the cytoarchitectonic borders against external landmarks is needed to develop the *in vivo* T1-weighted MRI protocol as the cytoarchitectonic borders of these cortical subregions are not visible on MRI. To develop rules for determining the anterior-most plane at which to begin labeling ERC, BA35 and BA36, for each of these regions we quantified the distance from the anterior-most histology slide that included the region of interest to various anatomical landmarks observable on MRI, including the temporal pole, collateral sulcus (CS), limen insulae (frontotemporal junction), amygdala, and the hippocampus. Supplementary Fig. 1 shows an example of each of the anatomical landmarks we assessed. These landmarks were assessed on histology sections except for the CS which was challenging to track on histology and was exclusively assessed on post-mortem MRI. Additionally, we used the post-mortem MRI to help identify the sulcal pattern for each case, as this is often more difficult to deduce based on histology slices.

To develop rules for the placement of ERC, BA35, and BA36 boundaries in the coronal plane, the distance from the border of interest to different landmarks observable on MRI were quantified in each histology slide using the Adobe Illustrator 2022 curvature tool (Supplementary Fig. 2). The depth of the CS was also measured to assess whether borders differed depending on the depth of the CS. A cut-off of 7mm in the first slide where the hippocampal head appears was used to determine if cases had a shallow or deep CS (28). 10 cases had a deep CS and 10 cases a shallow CS. The selection of landmarks to quantify their distances from each border was based on an initial qualitative assessment of the rough location of the border of interest on histology sections, e.g. it is observed that a specific border is in most cases on the crown of the parahippocampal gyrus.

Supplementary Fig. 3 shows landmarks we considered on histology and MRI.

### 2.4. ASHS-T1 and atlas set

The Automatic Segmentation of Hippocampal Subfields (ASHS) pipeline uses the combination of multi-atlas label fusion and corrective machine learning techniques (17,29) to map expert-generated segmentations from a set of example MRI scans, called atlases, to new, unlabeled MRI scans. The original ASHS pipeline requires high-resolution oblique coronal T2-weighted MRI scans, in which hippocampal layers are typically visible (17). However, since the segmentation of MTL cortical subregions such as ERC, BA35 and BA36 does not require hippocampal layer visibility, and since dedicated imaging of hippocampal subfields using T2-weighted MRI is not acquired in most studies, Xie et al. (1) adapted ASHS to 1x1x1mm^3^ T1-weighted MRI by transferring anatomical labels from the T2-weighted MRI to the same-subjects T1-weighted MRI in the ASHS atlas set (1). The ASHS-T1 pipeline segments the extrahippocampal MTL regions, but does not segment separate hippocampal subfields due to lack of contrast between hippocampal layers on most T1-weighted MRI scans (1). The ASHS atlas set consists of 15 cognitively normal older controls (NC) and 14 amnestic mild cognitive impairment (aMCI) patients, diagnosed according to established criteria (30–32). Demographic data for the aMCI and NC groups are shown in Supplementary Table 2.

The current ASHS segmentation protocol includes parahippocampal cortex, ERC and PRC (subdivided into BA35 and BA36) segmentations, however, these are limited to 1.3 mm anterior to the hippocampal head. The ASHS-T1 atlas also includes the anterior and posterior hippocampus (1) and has recently been extended to the amygdala (33). Our developed protocol will be applied to the ASHS-T1 atlas set to further expand the MTL cortex labels anteriorly.

### 2.5. ASHS-T1 atlas set image acquisition

MRI scans in the atlas set were obtained on a 3 T MRI scanner (Siemens Trio, Erlangen, Germany) at the University of Pennsylvania with an eight-channel array coil. A whole brain T1-weighted (magnetization prepared rapid acquisition gradient echo) MRI scan was obtained with Repetition Time (TR)/Time to Echo (TE)/Inversion Time (TI) = 1600/3.87/950 ms, 15^°^ flip angle, 1.0 × 1.0 × 1.0 mm^3^ isotropic resolution, and 5:13 min acquisition time. Additionally, a T2-weighted (turbo spin echo) MRI scan with partial brain coverage and oblique coronal slice positioned orthogonally to the main axis of the hippocampus (34,35) was obtained with TR/TE = 5310/68 ms, 18.3 ms echo spacing, 15 echo train length, 150^°^ flip angle, 0% phase oversampling, 0.4 × 0.4 mm^2^ in-plane resolution, 2.0 mm slice thickness with 0.6 mm gap, 30 interleaved slices, and 7:12 min acquisition time.

To reduce discrepancies in image resolution, the T1-MRI was upsampled to 0.5x0.5 in-plane resolution in the coronal plane using a patch-based super-resolution technique (36), and the T2-MRI was upsampled to 1.3 mm slice thickness. Affine registration between the upsampled T1-MRI and T2-MRI was performed separately in the left and right MTL region to best account for head motion between scans. The T1-MRI was then resliced into the space of the upsampled T2-MRI (0.4x0.4x1.3 mm3 resolution). Subsequently, anatomical labels of the original protocol (not including anterior labels for ERC, BA35 and BA36) from the T2-based ASHS atlas were transferred into the resliced T1-MRI, and manual corrections were made to account for remaining registration errors. A label for dura mater was added in the T1-MRI since dura matter and gray matter have similar appearance in T1-MRI (1).

### 2.6. Manual segmentation, reliability analysis, and cross-validation

After the development of the protocol, rater S.L. was trained on MR images that were similar to the atlas set, i.e. including both the high-resolution T2- weighted and the super-resolution T1-weighted images. Segmentations were performed on the super-resolution T1-weighted images, but the T2-weighted images were used to help infer grey/white matter boundaries in anterior regions where there are greater partial volume effects due to greater cortical curvature. The segmentations were performed using the ITK-SNAP image segmentation tool, version 4.0 (37). Following the training period, anterior ERC, BA35, and BA36 regions were segmented on 15 randomly selected cases of the ASHS atlas set by the rater, blinded to diagnosis. After a two-week interval, the protocol was applied to the same 15 cases but in a different order. The Dice Similarity Index (DSI) was calculated for each region. Then the remaining 14 cases of the atlas set were segmented.

ASHS-T1 standard five-fold cross-validation was performed, as in prior ASHS papers(38). The set of subjects with manual segmentations (n=29) was divided into five groups at random. Five experiments were performed. In each experiment, one group was held out for testing, and the remaining groups were used for training an ASHS model. DSC between automatic segmentations obtained using this trained model and the manual segmentations was computed for each subject in the held-out group. DSC values were averaged across all subjects in the held-out group and across the five experiments.

## 3. Results

### 3.1. Anterior boundaries of ERC, BA35, BA36

The distances between anterior histological borders of ERC, BA35 and BA36 and various anatomical landmarks observable on MRI were measured, including the temporal pole, collateral sulcus, limen insulae, amygdala, and the hippocampus. Supplementary Fig. 1 provides information on the detection of these landmarks on histology. Note that external landmarks are needed as the anterior borders of these cortical subregions are not visible on MRI.

Next, we compared the distances between the anterior borders of the MTL cortical subregions and various landmarks to determine the most suitable anchor point of these borders on MRI. Measured distances between MTL subregions and two of the landmarks, the anterior tip of the hippocampus and the anterior tip of the amygdala, had consistently lower standard deviations across the histology dataset than the distances to the remaining three landmarks (Table 2). Although the amygdala had a slightly lower between-subject variability compared to the hippocampus, the anterior tip of the amygdala is more challenging to identify on MRI images than the anterior limit of the hippocampus. Accordingly, we picked the anterior tip of the hippocampus as the most suitable anchor point.

**Table 2.** The distance (mm) of the MTL cortical regions to different landmarks observable on MRI. Bolded values are of the landmarks with the lowest standard deviations for the distances, suggestive of lowest between-subject variability. *Distances were measured on histology except for the collateral sulcus which was measured on MRI as it could not be identified reliably on the histology sections. Abbreviations: SD: standard deviation; ERC: entorhinal cortex; BA: Brodmann area

|  | ERC |  |  | BA35 |  |  | BA36 |  |  |
| --- | --- | --- | --- | --- | --- | --- | --- | --- | --- |
| **Distance\***<br>**from anterior**<br>**tip of:** | **Mean** | **Median** | **SD** | **Mean** | **Median** | **SD** | **Mean** | **Median** | **SD** |
| Hippocampus | **4.38** | **4.75** | **1.58** | **9.75** | **9.25** | **2.60** | **10.93** | **10.25** | **3.84** |
| Amygdala | **0.80** | **1.25** | **1.46** | **6.18** | **5.75** | **2.47** | **7.35** | **7.50** | **3.83** |
| Temporal pole | 21.28 | 21.75 | 2.22 | 15.90 | 15.50 | 3.27 | 14.73 | 15.00 | 3.81 |
| Limen insulae | 0.98 | 0.75 | 1.93 | -4.40 | -4.00 | 2.53 | -5.58 | -5.25 | 3.88 |
| Collateral sulcus | 15.07 | 14.80 | 4.36 | 9.69 | 9.20 | 4.43 | 8.69 | 7.05 | 5.28 |

Based on median values of measured distances, the ERC starts 4.75 mm, BA35 9.25 mm and BA36 10.25 mm anterior to the hippocampus, marking the location on the coronal plane to start segmenting the desired subregions. As our resampled *in vivo* T1-weighted MRI images have a slice thickness of 1.3 mm, we will develop a protocol for ERC starting 5.2 mm anterior to the hippocampus, for BA35 9.1 mm and for BA36 10.4 mm anterior to the hippocampus. These distances can easily be adapted when the protocol is applied to images with a more typical 1 mm slice thickness. As the posterior part of these regions is already included in the ASHS atlas, the posterior border did not need to be determined based on histology. The segmentation of these regions in the last version of the atlas starts at 1.3 mm anterior to the hippocampal head.

### 3.2. Developing segmentation rules

Due to the impracticality of performing the measurements on all histology slides available for each participant, we chose a set of 6 slices spaced throughout the length of the MTL cortex for development of segmentation rules, between 1 mm and 10 mm anterior to the hippocampus.

Slices at approximately the following locations were selected:

1. 10 mm anterior to the hippocampus (which is, on average, the anterior border of BA36)
2. 9 mm anterior to the hippocampus (which is, on average, the anterior- border of BA35)
3. 7 mm anterior to the hippocampus
4. 5 mm anterior to the hippocampus (which is, on average, the anterior border of the ERC)
5. 4 mm anterior to the hippocampus (which is, on average the anterior border of the amygdala)
6. 2.5 mm anterior to the hippocampus

Fig. 2 shows the sections selected relative to amygdala and the hippocampus.

**Figure 2.**
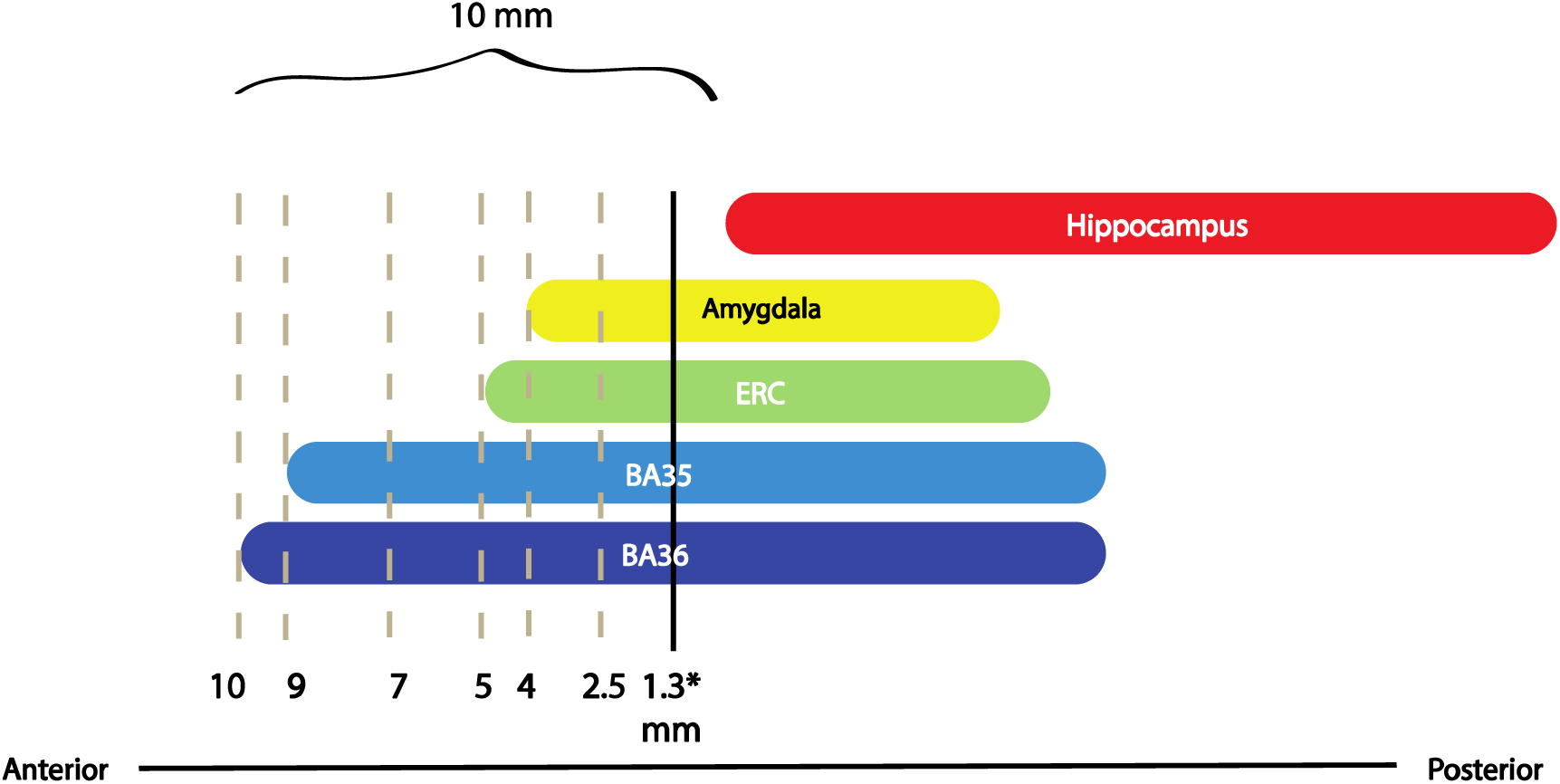
Histology sections selected for the rule development of the ERC, BA35 and BA36 boundaries in the coronal plane. Dashed lines mark the locations we chose for examining the histology slices. *The black line indicates the anterior border of the current protocol of the segmentations included in ASHS-T1. Abbreviations: ERC: entorhinal cortex; BA: Brodmann area

### 3.2. ERC boundaries

In our initial qualitative assessment of the general location of ERC borders we observed a consistent pattern; the medial border emerged roughly at the midpoint of the crown of the PHG in the first slice, then promptly shift to superior edge of the PHG (See Supplementary Fig. 3) as it progresses posteriorly, consistently maintaining this position. The lateral border was consistently located around the vicinity of the medial edge of the CS (Supplementary Fig. 3). Note that while the medial borders are located more superiorly, and the lateral borders are more inferiorly positioned, we will refer to all borders with “medial” and “lateral” throughout this manuscript to ensure consistency between the different sections.

We used median values to compare the distances and determine the borders.

Our measurements confirmed our initial assessments (Table 3). We showed that irrespective of the depth of the CS, the medial border corresponded to the superior edge of the PHG, while the lateral border of the ERC consistently aligned with the medial edge of the CS. Our findings for the medial border were consistent among all of the sections we inspected, except the most anterior slice. The most anterior section on which we investigated ERC borders was located 5 mm anterior to the hippocampus. However, in eight of the cases, the ERC was not yet present at that location, while in the remaining 12 cases, the SD for the medial border of the ERC in that section was relatively high (=5.06). This prompted us to instead examine anterior-most slide where ERC is present in all of our cases (this slice does not correspond to 5 mm anterior to the head of the hippocampus in all of the cases). We measured the distance from the medial border to both the superior edge of the PHG and to the halfway point of the crown of the PHG. Comparing the medians and SDs of the two measurements validated our initial qualitative assessment that the segmentation rule for the medial border of ERC on the anterior-most slice was more appropriately placed at the midpoint of the crown of the PHG (Fig. 3 and Table 6).

**Figure 3.**
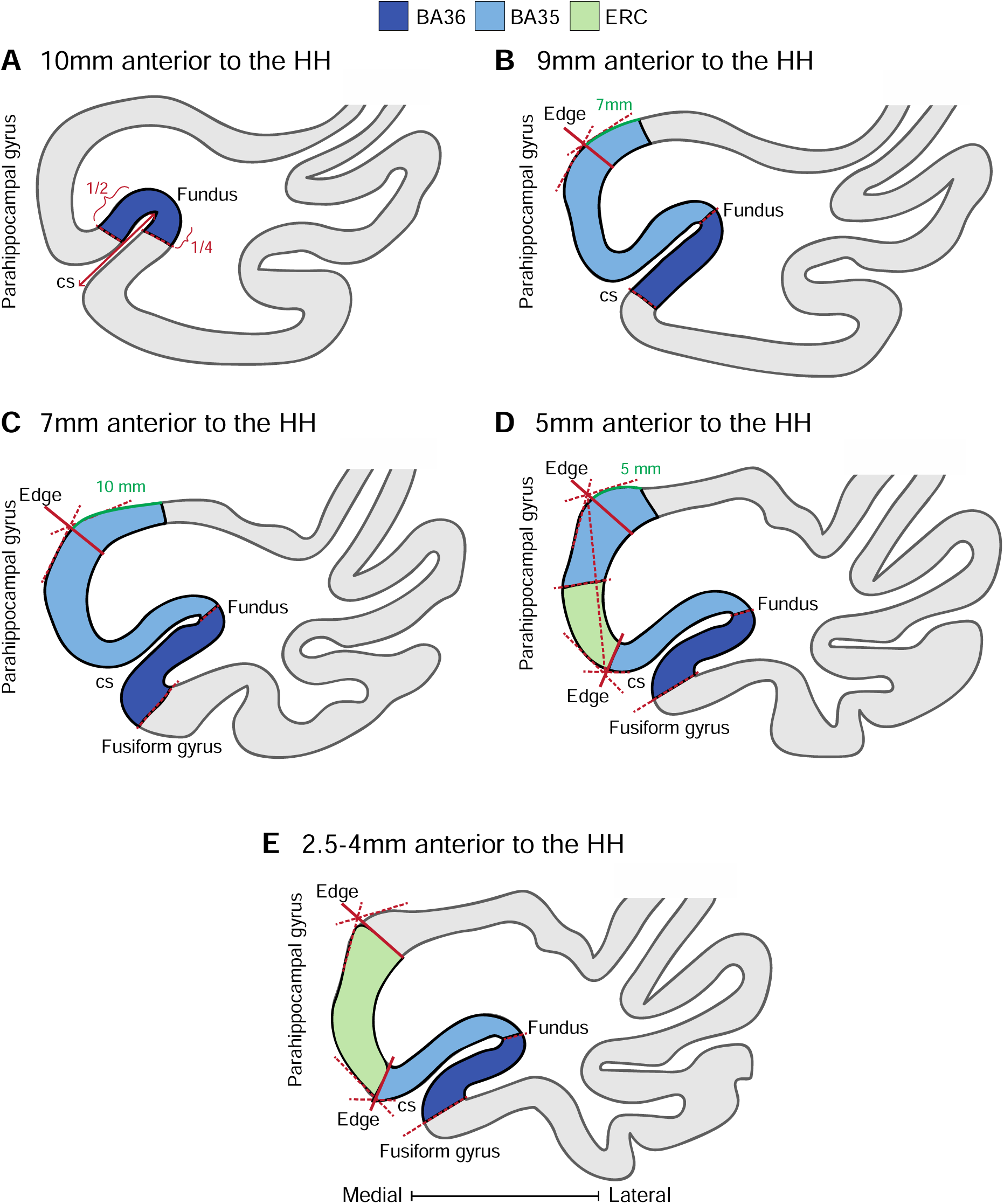
Application of the protocol on histology slides 10 mm to 2.5 mm anterior to the hippocampal head. Abbreviations: ERC: entorhinal cortex; BA35: Brodmann area35; BA36: Brodmann area36; CS: collateral sulcus; HH: hippocampal head

**Table 3.** Measured distances from the cytoarchitectonic borders of ERC to the chosen landmarks. Cells highlighted in grey were not used to formulate rules. For all borders, a negative value reflects the situation where the actual border is located medial of the chosen landmark and a positive value reflects the situation where the actual border is located lateral to the chosen landmark. * 5 mm anterior to hippocampal head, ERC had not appeared in 8 of the cases yet. Going more posterior to 4 mm anterior to the hippocampus, the ERC had not appeared in only one case. Abbreviations: ERC: entorhinal cortex; CS: collateral sulcus; PHG: parahippocampal gyrus; SD: standard deviation; H: hippocampus

| **From**<br>**To** | **Lateral ERC border** |  |  | **Medial ERC border** |  |  |  |  |  |
| --- | --- | --- | --- | --- | --- | --- | --- | --- | --- |
|  | **Medial CS edge** |  |  | **Superior PHG edge** |  |  | **Halfway point of PHG crown** |  |  |
| **Selected**<br>**slide:** | **Mean** | **Median** | **SD** | **Mean** | **Median** | **SD** | **Mean** | **Median** | **SD** |
| 5mm anterior to H (n=12) * | -0.05 | -1.85 | 4.14 | -2.97 | -2.38 | 5.06 | 5.55 | 6.88 | 6.13 |
| First ERC slide (n=20) | 0.75 | -0.83 | **4.39** | -5.08 | -4.50 | 6.24 | 1.17 | 0.92 | **4.20** |
| 4mm anterior to H (n=19) * | -0.62 | -0.88 | **3.45** | -2.60 | -3.11 | **4.02** |  |  |  |
| 2.5mm anterior to H (n=20) | -1.75 | -2.44 | **2.59** | -0.52 | -1.33 | **3.48** |  |  |  |

### 3.2. BA35 boundaries

Our initial inspection for BA35 borders for the full length of BA35 showed that the medial border was approximately at the superior edge of the PHG. However, we noticed a specific pattern for the more anterior sections in which ERC first becomes visible as well. In these sections, the ERC was embedded between two segments of BA35. On average this pattern was present for the first 2 histology sections of ERC with 0.5 mm distance (approximately 1 mm). This was, on average, 5 mm anterior to the hippocampal head. Based on our measurements (Table 4), for the anterior-most slice of BA35 located 9 mm anterior to the hippocampus, the medial border of BA35 is located 7 mm medial to the superior PHG edge. However, when determining the borders for the most anterior slice of BA35 we observed that in 6 cases BA35 had not yet appeared 9 mm anterior to the hippocampal head. As a result, we decided to also measure the borders on the most anterior slice where BA35 becomes visible in each case. The same border location was found when considering the measurements on the first slide of BA35.

**Table 4.** Measured distances from the cytoarchitectonic borders of BA35 to the chosen landmarks. For all borders, a negative value reflects the situation where the actual border is located medial of the chosen landmark and a positive value reflects the situation where the actual border is located lateral to the chosen landmark. *Not measured as this border is the ERC. ** The missing cases were due to either the region of interest not being present yet or deviant anatomy in that section that prevented us from doing the measurements. Abbreviations: BA35: Brodmann area 35; CS: collateral sulcus; PHG: parahippocampal gyrus; SD: standard deviation; H: hippocampus

| **From** | **Lateral border of BA35** |  |  | **Medial border of BA35** |  |  |
| --- | --- | --- | --- | --- | --- | --- |
| **To** | **CS fundus** |  |  | **Superior PHG edge** |  |  |
| **Selected slide:** | **Mean** | **Median** | **SD** | **Mean** | **Median** | **SD** |
| 9mm anterior to H<br>(n=14) ** | -0.52 | 0.00 | 4.86 | 8.15 | 7.19 | 4.35 |
| First BA35 slide<br>(n=17) ** | -4.21 | -1.89 | 6.99 | 0.84 | 6.03 | 11.54 |
| 7mm anterior to H<br>(n=17) ** | -1.77 | 0.00 | 5.01 | 10.44 | 9.96 | 4.10 |
| 5mm anterior to H<br>(n=16) ** | 0.00 | 0.30 | 2.32 | -6.18 | -5.40 | 3.15 |
| 4mm anterior to H<br>(n=19) ** | 0.46 | 0.86 | 2.11 | N/A* |  |  |
| 2.5mm anterior to H<br>(n=20) | 0.55 | 0.81 | 2.13 | N/A* |  |  |

Going more posterior to the slice located 7 mm anterior to the hippocampus, the border shifts to be located 10 mm medial to the superior PHG edge. When ERC becomes visible 5 mm anterior to the hippocampal head, there are two segments of BA35 surrounding ERC. The most medial border of BA35 here is located 5 mm medial to the superior PHG edge. For the rest of the slides located more posteriorly (1-5 mm anterior to the hippocampus) the lateral border of ERC serves as the medial border of BA35. The lateral border is in the CS fundus for the full length of BA35, including the second section of BA35 lateral to the ERC where ERC first emerges (Fig. 3 and Table 6).

### 3.2. BA36 boundaries

The qualitative evaluation of BA36 borders for the full length of this region showed that the medial border was approximately located around the medial bank of the CS anteriorly and gradually moved to the fundus of the CS towards the posterior end. The lateral border was anteriorly detected close to the fundus of the CS. Moving posteriorly, the lateral border changed gradually towards the crown of the fusiform gyrus (FG).

Based on our measurements (Table 5), the medial border of BA36, 10 mm anterior to the hippocampal head, is located at the halfway point of the medial bank of the CS. However, when determining the borders for the most anterior portion of BA36, we noticed that in 5 of the cases BA36 had not emerged yet 10 mm anterior to the hippocampal head, and the standard deviation for the measurements were quite high at this level. Therefore, we decided to measure the borders on the most anterior slide where BA36 becomes visible in each case. The same border location was found when considering the measurements on the first slide of BA36. For all remaining slices from 1-9 mm anterior to the hippocampus, the lateral border of BA35 serves as the medial border of BA36.

**Table 5.** Measured distances from the cytoarchitectonic borders of BA36 to the chosen landmarks. Cells highlighted in grey were not used to formulate rules. For all borders, a negative value reflects the situation where the actual border is located to the medial of the chosen landmark and a positive value reflects the situation where the actual border is located lateral to the chosen landmark. *Not measured as the border is BA35. ** The missing cases were due to either the region of interest not being present yet or the deviant anatomy in that section that prevented us from doing the measurements. Abbreviations: BA36: Brodmann area 36; CS: collateral sulcus; SD: standard deviation; H: hippocampus; FG: fusiform gyrus

| From | Lateral border of BA36 |  |  |  |  |  | Medial border of BA36 |  |  |
| --- | --- | --- | --- | --- | --- | --- | --- | --- | --- |
| To | CS fundus |  |  | Halfway point of the lateral bank of the CS |  |  | Halfway point of the medial bank of the CS |  |  |
| Selected slide: | Mean | Median | SD | Mean | Median | SD | Mean | Median | SD |
| 10mm anterior to H (n=15) ** | 8.07 | 7.67 | 6.21 | 3.73 | 1.86 | 7.39 | -2.85 | 0.00 | 7.27 |
| First BA36 slide (n=18) ** | 3.34 | 0.5 | 5.37 | 0.58 | -1.07 | 6.28 | -0.61 | 2.02 | 7.94 |
| To | Halfway point of the crown of the FG |  |  |  |  |  |  |  |  |
|  | Mean |  | Median |  | SD |  |  |  |  |
| 9mm anterior to H (n=11) ** | 0.66 |  | -0.62 |  | 8.69 |  | N/A* |  |  |
| 7mm anterior to H (n=15) ** | -2.75 |  | -2.36 |  | 6.91 |  | N/A* |  |  |
| 5mm anterior to H (n=18) ** | -0.63 |  | 0.00 |  | 7.71 |  | N/A* |  |  |
| 4mm anterior to H (n=16) ** | 0.75 |  | 0.00 |  | 4.71 |  | N/A* |  |  |
| 2.5mm anterior to H (n=18) ** | 0.34 |  | 0.00 |  | 6.71 |  | N/A* |  |  |

**Table 6.**
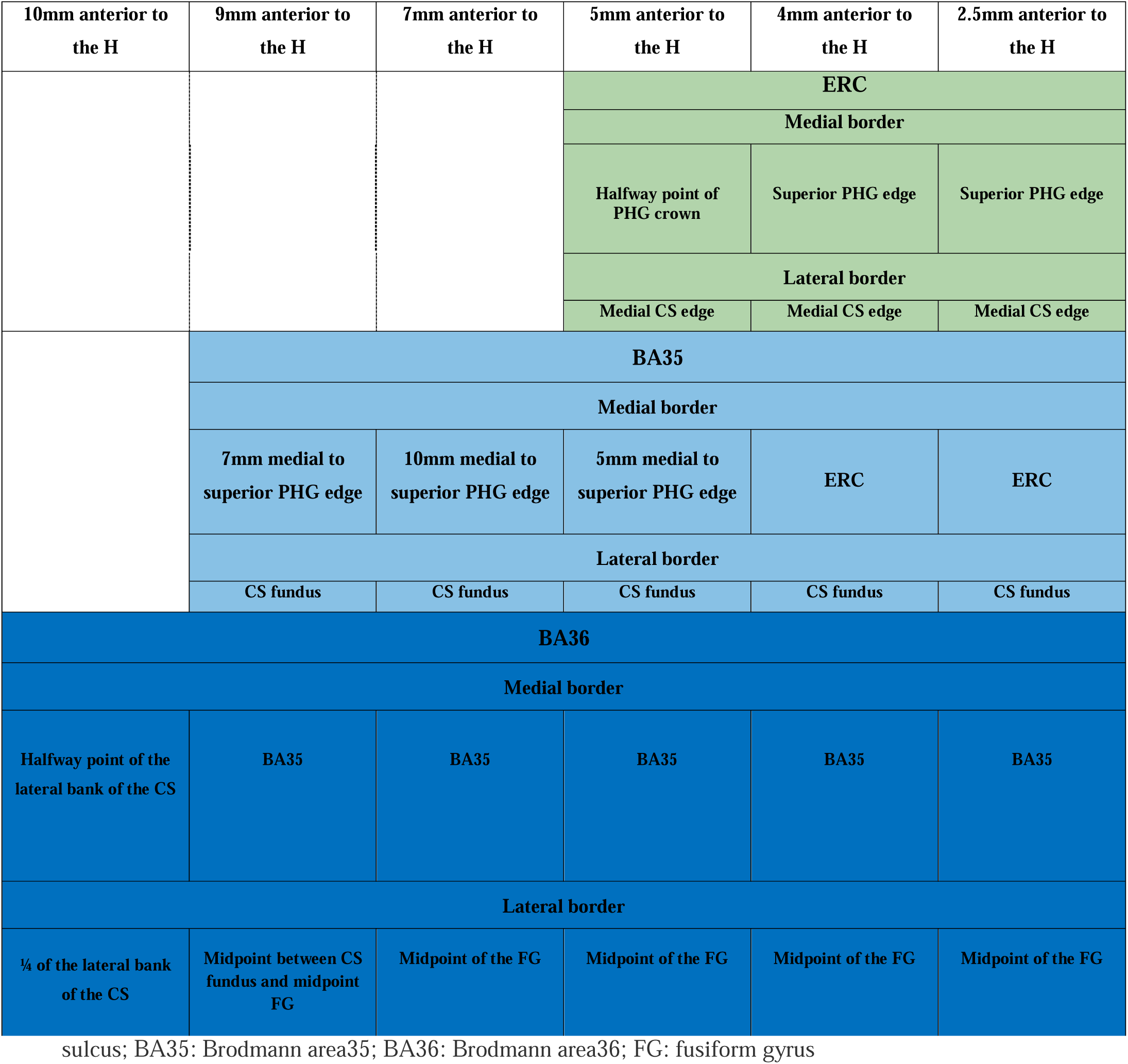
Overview of the rules for border placement for the entorhinal cortex, Brodmann area 35 (BA35), and BA36 in anterior medial temporal lobe. Abbreviations: H: the hippocampus; ERC: entorhinal cortex; PHG: parahippocampal gyrus; CS: collateral

| 10mm anterior to the H | 9mm anterior to the H | 7mm anterior to the H | 5mm anterior to the H | 4mm anterior to the H | 2.5mm anterior to the H |
| --- | --- | --- | --- | --- | --- |
|  |  |  | **ERC** |  |  |
|  |  |  | **Medial border** |  |  |
|  |  |  | Halfway point of PHG crown | Superior PHG edge | Superior PHG edge |
|  |  |  | **Lateral border** |  |  |
|  |  |  | Medial CS edge | Medial CS edge | Medial CS edge |
|  | **BA35** |  |  |  |  |
|  | **Medial border** |  |  |  |  |
|  | 7mm medial to superior PHG edge | 10mm medial to superior PHG edge | 5mm medial to superior PHG edge | ERC | ERC |
|  | **Lateral border** |  |  |  |  |
|  | CS fundus | CS fundus | CS fundus | CS fundus | CS fundus |
| **BA36** |  |  |  |  |  |
| **Medial border** |  |  |  |  |  |
| Halfway point of the lateral bank of the CS | BA35 | BA35 | BA35 | BA35 | BA35 |
| **Lateral border** |  |  |  |  |  |
| ¼ of the lateral bank of the CS | Midpoint between CS fundus and midpoint FG | Midpoint of the FG | Midpoint of the FG | Midpoint of the FG | Midpoint of the FG |

In the most anterior BA36 slide, the best location for the lateral border was the fundus based on our measurements. However, when looking at this border 10 mm anterior to the head of the hippocampus, the best place to put the medial border was determined to be the halfway point of the lateral bank of the CS. To take this difference into account, we decided to place this border at 1/4^th^ of the lateral bank of the CS- closest to the fundus of the CS. In more posterior slices (1- 9 mm anterior to the hippocampal head) the border shifts to be placed at the midpoint of the crown of the FG (Fig. 3 and Table 6). However, to keep the transition of the lateral border smooth, we determined this border to be at the midpoint between the fundus of the CS and midpoint of the crown of the FG for the second slice of BA36 located 9 mm anterior to the hippocampus.

Supplementary Fig. 4 depicts this modification in border placement.

### 3.3 Effect of the depth of the collateral sulcus and diagnostic status on proposed boundaries

Supplementary Table 3-5 shows the comparison of the borders’ locations when cases are divided based on the depth of the CS (deep vs. shallow) and between cases with neurodegenerative diseases vs. cases without neurodegenerative diseases. While some small differences were observed, there was not a consistent difference for these groups that warranted creating additional rules for the protocol. Moreover, differences in the SD can be observed between cases with a deep vs a shallow CS for some of the borders for BA35 and BA36, where a higher SD indicates larger variability in the border location between cases of that group. However, as the opposite pattern is observed for BA35 (SD lateral border 7 mm anterior to the hippocampus: deep>shallow)) and BA36 (SD lateral and medial border 10 mm anterior to the hippocampus: shallow>deep), it is unclear how this should be interpreted.

### 3.4. Inter-rater reliability and cross-validation evaluations in the atlas set in the space of T1w MRI (ASHS-T1)

Table 7 shows intra-rater reliability of the rater (S.L.) for 15 cases of the ASHS atlas set. All DSI values were above 0.8 and indicated high reliability.

**Table 7.** Intra-rater reliability of a single rater (S.L.) and ASHS-T1 cross validation for anterior MTL, whole MTL, and MTL without the new anterior region Cross-validation was performed separately for anterior slices of the MTL and for the whole MTL, including the more posterior slices which already existed in the original ASHS-T1. Abbreviations: MTL: medial temporal lobe; SD: standard deviation; ERC: entorhinal cortex; BA35: Brodmann area 35; BA36: Brodmann area 36

|  | Intra-rater reliability |  | ASHS-T1 cross-validation<br>(anterior MTL) |  | ASHS-T1 cross-validation<br>(whole MTL) |  | ASHS-T1 cross-validation<br>(MTL without the new anterior region)(1) |
| --- | --- | --- | --- | --- | --- | --- | --- |
| DSI (Mean±SD) |  |  |  |  |  |  |  |
| Region | Left | Right | Left | Right | Left | Right | Average |
| ERC | 0.88±0.02 | 0.86±0.05 | 0.64±0.12 | 0.62±0.12 | 0.77±0.04 | 0.75±0.05 | 0.76±0.03 |
| BA35 | 0.88±0.07 | 0.88±0.03 | 0.64±0.11 | 0.62±0.10 | 0.69±0.05 | 0.67±0.06 | 0.71±0.06 |
| BA36 | 0.86±0.08 | 0.85±0.08 | 0.65±0.11 | 0.63±0.08 | 0.78±0.04 | 0.78±0.04 | 0.79±0.03 |

To compare the segmentation accuracy of ASHS-T1 for the newly added anterior MTL cortices, cross-validation evaluation was performed. Table 7 shows the ASHS five-fold cross-validation results. The cross-validation results showed lower variability with DSIs in the range from 0.62 to 0.65 for the anterior MTL structures. The cross-validation DSI value were higher (0.67 to 0.78, indicating moderate reliability) when combining the anterior and posterior portions of the ERC, BA35 and BA36 labels. Indeed, DSI for the combined anterior/posterior labels was on par with previously reported cross-validation DSI for ERC, BA35, and BA36 without the new anterior extension (1). Fig. 4 compares a manual and an automated segmentation on one of the cases of our atlas set.

**Figure 4.**
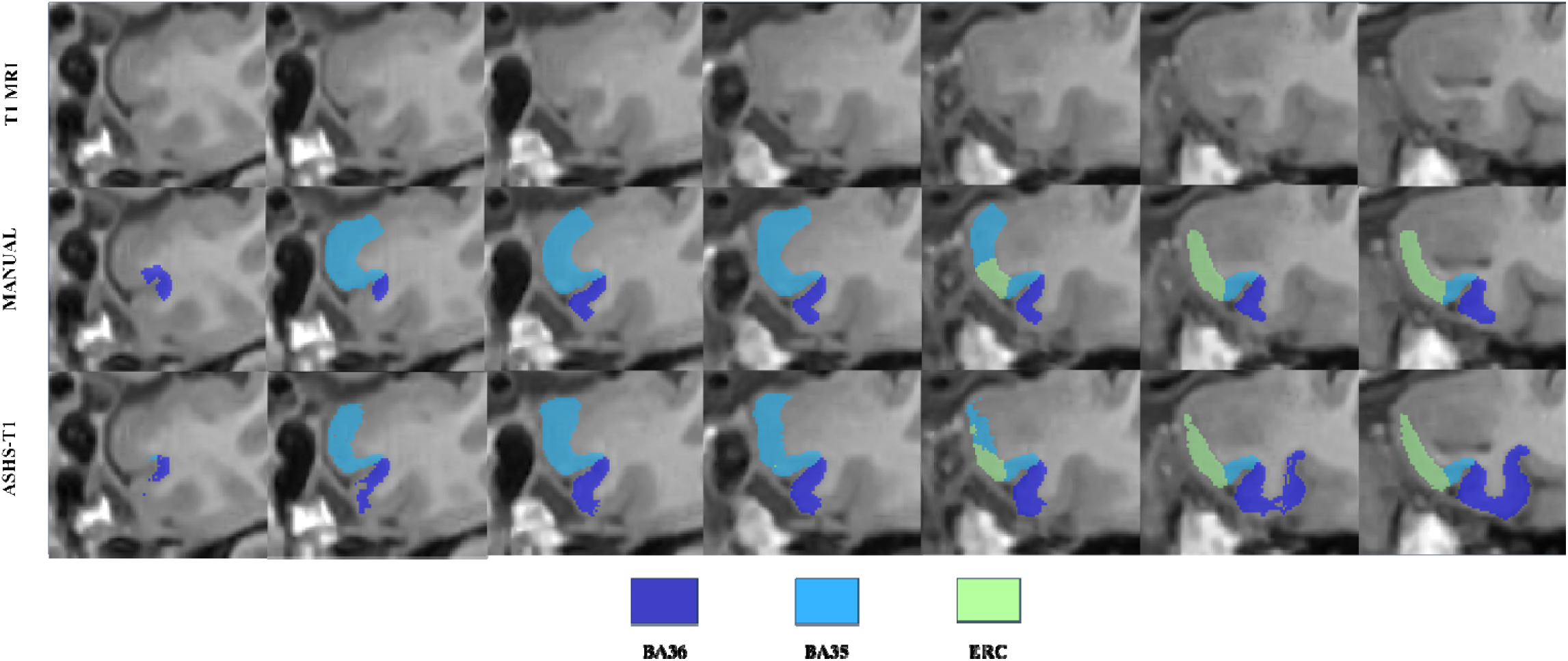
Comparing manual vs. automatic segmentation (ASHS-T1) on one of the cases of our atlas set. Abbreviations: ERC: entorhinal cortex; BA: Brodmann area

Supplementary Fig. 6 Shows 3D rendering of the new anterior regions for two cases from the ASHS atlas set, including one control subject and one individual with MCI. These 3D renderings show consistency of the borders between anterior MTL cortices between consecutive slices. Additionally, Supplementary Fig. 7 shows 3D rendering of the automatic segmentation of the anterior regions of the MTL for four cases from the ASHS atlas set, including two control subjects and two individuals with MCI.

Finally, we compared MTL cortex volumes between cases with a shallow versus a deep sulcus (Supplementary Table 6) to investigate whether sulcal depth has a significant effect on MTL cortex volumes. We found no evidence for this hypothesis.

## 4. Discussion

To the best of our knowledge, the present study is the first to present a manual segmentation protocol for anterior regions of the MTL grounded in a large set of cases with annotated histological sections. Moreover, the implementation on *in vivo* MRI showed high reliability for a manual rater and moderate reliability for an automated approach, when the whole MTL was taken into account. Both through the inclusion of 20 cases with varying anatomy and through the multi-atlas approach, we ensured that our approach is generalizable to a wide range of anatomical variability and to cases with and without neurodegenerative diseases.

While atlases exist that are based on either postmortem MRI or histological annotations (19,39), these atlases do not provide labels for anterior MTL cortical regions, including BA35, nor do they utilize a multi-atlas approach that can be applied to individual MRI scans thereby taking anatomical variability into account.

We included 20 cases to develop this protocol and to translate two-dimensional histologically recognized regions to neuroanatomically valid borders applicable to MRI scans. Having the corresponding post-mortem MRI scans of our cases enabled us to track CS patterns and check for deviant anatomy for each case. Gross assessment of the evolution of individual borders across our histology dataset enabled us to choose a number of potential landmarks for each of the borders. We systematically measured border distances to those potential landmarks in slides selected at varying distances to the hippocampal head to ensure capture of changing patterns for each border. Ultimately, we selected landmarks that showed the least between-subject variations in border-landmark distances and are easily identifiable on MRI.

Surprisingly, examining the diagnosis status during border placement in our protocol revealed minimal effect of a wide range of common neurodegenerative diseases on proposed borders. While this is a novel observation, the number of cases is still relatively small and insufficient to assess differences across the different neurodegenerative disease diagnoses. This will be an interesting avenue for future research. Additionally, we demonstrated that proposed borders were not dependent on the depth of the CS. This is in contrast to what has previously been reported in other neuroanatomical references (14,40), although previous work did not specifically focus on anterior versus posterior segments of the MTL cortical region borders and how these potentially differentially depend on CS depth. It is possible that the dependency of MTL cortex borders on the CS depth differs from more anterior to more posterior regions. However, the differences between deep versus shallow CS groups was variable in an anterior-to-posterior direction (Supplementary Table 3-5). Additionally, even the largest observed differences were so small that they did not warrant a change in segmentation rules. Finally, volumes MTL cortical regions, obtained from the in vivo automated segmentations, were also not different between subjects with a shallow versus a deep CS. Future work should explore this further, also in other populations.

Besides anatomical validity, our protocol also showed high intra-rater reliability for a manual rater, with DSI values that fell within the range of previous manual segmentation protocols for MTL cortices (15,28,38). The DSI values comparing automated segmentations with manual segmentations for the current protocol ranged between 0.62 and 0.65, which is relatively low, compared to previous studies (1,41,42). However, the reliability was higher when considering the combined (anterior and posterior) labels of the MTL cortices and more consistent with the reliability of these structures without the new anterior extension reported previously on the same atlas set (1), as well as within the range of DSI scores for MTL cortices reported in other datasets (1,41–43). The reason for the relatively lower DSI scores for the anterior segment can likely be attributed at least in part to the smaller size of these regions, which is known to be penalized by the DSI scores (44), and the increasingly complicated anatomy in the anterior MTL. The DSI scores of the combined anterior/posterior MTL cortex labels are more important though, as the MTL cortices will likely be analyzed as one region rather than separated based on a relatively arbitrary border, such as 1 mm anterior to the hippocampus. Moreover, instead of volumes, it is also possible to analyze the median thickness obtained from these labels, which is likely also less affected by errors in the automated segmentation.

While we developed an anatomically valid and reliable protocol for the anterior MTL, it should be noted that a limitation of this study is that the annotations of the histological sections were only performed by a single neuroanatomist. Recent work from the Hippocampal Subfields Group showed an in-depth characterization on border definitions and placement of MTL cortical regions of different neuroanatomy laboratories (45), where most disagreement was observed in transition zones between different regions, including the anterior-most point of the MTL. Our work is therefore limited by being based on the anatomical definitions of only one neuroanatomy laboratory. However, as the harmonized protocol for MTL cortical regions for T2-MRI of the Hippocampal Subfields Group is still some time away, and especially a potential adaptation to T1-MRI, we believe that the current expansion of ASHS will be a useful tool for the neuroimaging community in the meantime. Moreover, in our work we were able to analyze the anterior MTL in a relatively large set of histological annotations of 20 cases, which was not possible for the work of the Hippocampal Subfields Group because of the labor intensiveness of annotating such a large sample of cases.

Another limitation is that we were not able to assess the effect of intracranial volume on the border locations because of the relatively small sample size and the fact this this information is not available in a portion of our postmortem dataset.

While this is a limitation of all current segmentation protocols to the best of my knowledge, it will be interesting to explore the effect of intracranial volume on border locations and perhaps anatomical landmarks. A final limitation is that the protocol is based on histological sections from older individuals (61-97 years) and is implemented on MRI scans of an in vivo population consisting of older adults. This may limit the utility of our manual segmentation protocol and the application of our automated approach in younger or otherwise different populations.

## 5. Conclusion

There is increasing evidence suggesting that different diseases affect extrahippocampal cortices in the MTL such as the ERC, BA35 and BA36 and especially in anterior regions (22,24,46,47). The development and implementation of reliable and anatomically valid anterior MTL labels in our automated ASHS-T1 pipeline based on widely available T1-MRI, presented here, will enable the investigation of these sensitive MTL cortical regions in AD and LATE and hopefully lead to more sensitive biomarkers for early disease detection, monitoring, and clinical trials. This extension of ASHS-T1 will be made publicly available with this publication.

## Acknowledgment

We gratefully acknowledge the tissue donors and their families. We also thank all the staff at the Center for Neurodegenerative Disease Research (University of Pennsylvania) and Human Neuroanatomy Lab (University of Castilla-La Mancha—UCLM) for performing the autopsies and making the tissue available for this project.

