## supplementary_file for "Developing an anatomically valid segmentation protocol for anterior regions of the medial temporal lobe for neurodegenerative diseases"

**Supplementary Methods:
Criteria for medial temporal lobe subregion annotations on histological sections**

- **Introduction**

The annotation procedure has been systematically used throughout the series of cases irrespective of the origin of the cases.

For each case, the whole temporal lobe has been divided into 4 blocks, each one being 2 cm thick. 50 µm serial sections were obtained throughout every block, from beginning to end, which resulted approximately in 400 sections per block. The sections are processed in groups of 10 sections, also called series, resulting in a 500 µm distance among adjacent series. Of the series, 5 sections were put together in a freezing protectant solution in 24 wells plastic plaques. The following 4 sections were placed individually in labeled Eppendorf 1.5 cc tubes with cryoprotectant: retaining the order of the sections. Section number 10 was immediately mounted for Nissl staining. Blockface photographs after each section were obtained and stored. After processing for Nissl stain and other determinations, the resulting slides were scanned. The scanned sections were used for annotating, although slides were also examined under the microscope when necessary.

As a consequence, we had for annotations one section every 0.5 mm (500 µm), and a total of about 150 histological slides per case.

- **Procedure for annotating the Entorhinal cortex**

The Entorhinal Cortex (ERC) shows a primitive lamination and is considered as Periallocortex (1). The basis for the annotations of the ERC is derived from a cytoarchitectonic study by Insausti et al. (2), with additional information extracted from previous studies (1,3–7). The partition of the ERC in eight subfields follows the general framework of former anatomists who considered the ERC as a heterogeneous region, and defined differences in layering. Here, we followed the scheme of lamination of Cajal (8), in particular for layer IV (*Lamina dissecans*), while keeping the number of layers described by Lorente de Nó (5). Based on the observation of that the structure and relative thickness of each layer differed per ERC subfield, the annotations followed features specific for each one of the subfields (1,2).

Moving in an anterior-to-posterior (rostral-to-caudal) direction, the first ERC subfield emerging close to the medial edge of the collateral sulcus, is the Entorhinal Lateral Rostral subfield (EL_R_). Simultaneously, or shortly thereafter, superior to EL_R_ and bordering the periamygdaloid cortex, the Entorhinal Olfactory subfield (E_O_) shows up. A few mm in the posterior direction lies the Entorhinal Rostral (E_R_) subfield, that shows a more organized lamination. Approximately at midlevel of the amygdaloid cortex E_R_ is replaced by the Entorhinal Intermediate subfield (E_I_), which covers almost the entire medial aspect of the parahippocampal gyrus, with EL_R_ bordering inferiorly. Further posteriorly, the Entorhinal Medial Intermediate (EMI) appears. This region is characterized by the clearest lamination of all subfields of the ERC. Additionally, E_MI_ predominantly occupies the Gyrus ambiens, which is a medial bulge located on the medial aspect of the MTL. At this level, EL_R_ is replaced by the Entorhinal Lateral Caudal subfield (EL_C_), which occupies the same topographical position as EL_R_, though generally located on the medial bank of the collateral sulcus. The emergence of the Entorhinal Caudal subfield (E_C_) is roughly indicated by the opening of the hippocampal fissure, marking the gradual replacement of the E_I_ and EL_C_ subfields. The Entorhinal Caudal Limiting subfield (ECL), which is the most posterior among the ERC subfields, begins either simultaneously or shortly after this point. ECL partially intermingles superiorly with the boundaries of Presubiculum or Parasubiculum, as well as with area TH of the parahippocampal cortex.

The annotation of all subfields of the ERC was conducted in accordance with this topographical organization and the previously mentioned distinctions in lamination.

- **The procedure for the annotation of the remaining medial temporal lobe cortices**

We took the junction of the frontal and temporal lobes (*Limen* *insulae* or frontotemporal junction, i.e. plate 26 in Mai et al. (9)) as the starting point for the annotation of cortical regions. This area is the intersection of several cytoarchitectonic fields, including the posterior end of the temporopolar cortex and the anterior-most limit of Brodmann Area (BA) 36 (ectorhinal cortex, later subsumed under the term perirhinal cortex (PRC), along with BA35). Lateral to the *Limen insulae* lies the inferior temporal cortex. The cortex between the collateral sulcus and the inferior temporal sulcus makes up the fusiform gyrus. Laterally from the fusiform gyrus, the inferior temporal gyrus is located, which corresponds to inferotemporal cortex or BA20 (also named area TE by von Bonin and Bailey (10)). The boundary between BA36 and area TE (BA20) lies along the surface of the fusiform gyrus. However, the exact location of this boundary is not precisely defined, as it depends on the shape of the MTL and its sulci, in particular the collateral sulcus.

The cortical areas annotated here were identified based on our own publications (11–13) along with other previously published descriptions and atlases (9,10,14–16).

The nomenclature of the temporopolar cortex (BA38) in the annotations follows Salinas (11), with extensions described by Blaizot et al. (12). Essentially, the temporopolar cortex (BA38) is smaller than the original delineation by Brodmann (3).

Proceeding anteriorly from the *Limen insulae*, we annotated the transition between temporopolar cortex and PRC (BA35+36), as well as those with area TE (inferotemporal cortex). Eventually, we reach a point where only the medial and lateral regions (TPCm – medial part, TPCl – lateral part) of the temporopolar cortex are present (9,11,12,17). Once the anterior part of the MTL is annotated, we continue to the posterior-most end of the MTL.

Several of the aforementioned cortical subfields (e.g. PRC with both BA35 and BA36 divisions, part of the posterior portion of the medial and lateral divisions of temporopolar cortex, and anterior area TE (inferotemporal cortex) are already present at this point. While the collateral sulcus is present here, it exhibits extensive morphological variability and offers little assistance in the annotations.

Moving a few mm in a posterior direction, it is possible to find the anterior-most part of the entorhinal cortex (ERC). The nomenclature employed in the annotations of the ERC subfields follows our own framework (2), which was also used by other authors (i.e. Mai et al.(9)). The first two subfields that usually emerge in the series of sections are the olfactory (E_O_) and laterorostral (EL_R_) subfields. Those ERC subfields are adjacent to BA35, and their presence limits BA35 to either the medial bank or the fundus of the collateral sulcus; the depth of the sulcus helps with orienting the boundary of ERC and BA35, as described by Insausti et al. (17). In a lateral direction lies BA36 along the lateral bank of the collateral sulcus and dorsomedial portion of the surface of the fusiform gyrus. The rostromedial portion of area TE is situated lateral to BA36. This boundary with BA36 is far from clearcut. Therefore, the annotations of this part of the MTL may vary among different subjects.

Further posteriorly more ERC subfields become visible, including the rostral (E_R_) and intermediate (E_I_) subfields. Their annotation follows the criteria described by Insausti et al. (2). The ERC and the medial-most portion of BA35 adopt a particular wedge shape, previously identified by several authors (9,11,14,15), compatible with the transentorhinal cortex described by Braak and Braak (18). The annotated boundary of this portion of BA35 can be located close to the fundus, on the midportion of the medial bank or close to the edge of the medial bank of the collateral sulcus. The typical emergence of the transentorhinal portion of BA35 is followed and complemented by another part of BA35 (35v - ventral) that is usually located at the fundus of the collateral sulcus and for a few mm, into the lateral bank of the collateral sulcus. Here, it borders the most medial part of BA36, but the boundary may be difficult to determine, as the transition from one to the other is gradual. This characteristic produces some variability in annotations among subjects.

BA36 extends from the 35v portion of BA35 into the surface of the brain, as it is usually present in the crown of the fusiform gyrus. The extent of the crown of the fusiform gyrus, along with the depth of the collateral sulcus determines the variability observed among subjects (19). Additionally, the transition between area TE and BA36 is very gradual, showing variability among subjects. However, typically, this boundary is situated around the crown of the fusiform gyrus or near the medial edge of the occipitotemporal sulcus, which is positioned laterally to the fusiform gyrus. We distinguished two divisions in BA36, one situated anteriorly (36r-rostral) and one further posteriorly (36c-caudal). Although both divisions maintain a consistent topographical relationship with adjacent fields (35v, TE) in terms of mediolateral extent, the organization of their constituent layers starts to resemble that of the neocortex at more posterior levels. The posterior end of 36c overlaps slightly with the anterior extreme of the parahippocampal cortex (PHC), as it shares some similarities in overall appearance. The structural organization of the cortical layers resembles the neocortex present in the inferotemporal cortex (TE). At this level of the brain, the annotations are heterogeneous because the confluent cortical fields may exhibit different extensions and shapes (19). We followed our own studies on the MTL (9,11), in which MTL annotations were made by RI. A representative illustration can be seen in plate 50 of Mai et al. (9). Additional information was obtained by Bailey and von Bonin (10).

Given the confluence of various cortical MTL areas in this specific place, and the resultant complexity, a useful macroscopic landmark, is indicated by the termination of the uncus, which is marked by the Gyrus intralimbicus. This structure is easily recognizable by its appearance as a rounded structure, either connected to the rest of the hippocampus by the fimbria as a thin white matter thread, or as a separate entity.

The posterior end of the entorhinal cortex marks the transition to the posterior part of the parahippocampal gyrus. The ECL subfield of the entorhinal cortex is continued by the anterior extreme of the PHC. We recognized two main parts in the PHC organization: area TH and area TF (based on von Economo and Koskinas (20) and followed by a study by Bonin and Bailey (10) in humans). Area TF has been subdivided in area TFm (medial), situated anteriorly, which overlaps with 36c, and area TFl (lateral), which extends to the end of the parahippocampal gyrus, at the level of the hippocampal tail. The posterior portion of TFl borders the anterior extreme of the retrosplenial cortex. Annotations of the PHC are more uniform at mid and posterior levels of the PHC, whereas at anterior levels, annotations must account for the confluence of multiple MTL areas or subfields. At a particular section, one may encounter the posterior end of the ERC (subfield ECL) along with the anterior extension of area TH. Simultaneously, area 36c of the PRC may be present, extending laterally. Additionally, this section might also contain the anterior extension of area TFm, partially covering the medial bank of the collateral sulcus, while the remaining bank of the collateral sulcus corresponds to area TE. This level of the MTL has high complexity, where many boundaries are present in one single section. We followed our own description of the cortex of the PHC (21), based on the basic criteria of von Economo and Koskinas (20) and Bailey and von Bonin (10) (1951). Similar definitions are followed in Mai et al. (9). It is worth noting that the PHC, although studied in nonhuman primates, is only scarcely studied in humans.

**Supplementary results:**

**Supplementary Table 1** Demographic and diagnostic details for subjects included in this study.

Abbreviations: ADNC: Alzheimer’s disease neuropathologic change; CVD: cerebrovascular disease; CAA: cerebral amyloid angiopathy; CBD: corticobasal degeneration; LBD: Lewy body disease; FTLD-PPA (PNFA): frontotemporal lobar degeneration-primary progressive aphasia (progressive nonfluent aphasia); PD: Parkinson’s disease; PSP: progressive supranuclear palsy; PART: primary age-related tauopathy; FTLD-bvFTD: frontotemporal lobar degeneration-behavioral variant frontotemporal dementia; FTLD-TDP: frontotemporal lobar degeneration with TDP-43 inclusions.

| **Patient ID** | **Hemisphere** | **Age (years)** | **Sex** | **Clinical Diagnosis** | **Neuropathological Diagnosis** | **PMI (hours)** |
| --- | --- | --- | --- | --- | --- | --- |
| HNL-01 | L | 78 | F | Unknown | Low ADNC, CVD | 6 |
| HNL-02 | R | 90+ | M | Unknown | Intermediate ADNC, CAA | 11 |
| HNL-03 | R | 74 | M | Unknown | Pathological aging (mild tau pathology, B2), and mild CAA | 3 |
| HNL-04 | R | 61 | M | Unknown | Pathological aging (mild tau pathology, B1), suspected CBD | 16 |
| HNL-05 | R | 76 | F | Unknown | No pathology | 4 |
| HNL-06 | L | 66 | F | Unknown | No pathology, Incidental LBD | 9 |
| HNL-07 | R | 90+ | F | Unknown | Low ADNC | 16 |
| HNL-08 | L | 90 | M | Unknown | Low ADNC, Brainstem predominant Incidental LBD | 7 |
| HNL-09 | R | 74 | M | Unknown | Low ADNC | 2 |
| HNL-10 | L | 62 | F | Unknown | Low ADNC | 3 |
| INDD-01 | R | 76 | F | FTLD-PPA (PNFA) | 1.CBD 2. low ADNC | 14 |
| INDD-02 | R | 90+ | M | PD with Dementia | 1.LBD, 2. high ADNC | 17 |
| INDD-03 | L | 86 | M | Probable AD | 1. high ADNC ,2. LATE | 10 |
| INDD-04 | R | 90+ | F | Normal | 1. intermediate ADNC, 2. LBD | 6 |
| INDD-05 | L | 79 | F | PSP | 1. PSP, 2. PART | 5 |
| INDD-06 | R | 70 | M | PD with Dementia | 1. LBD, 2. PART | 9.5 |
| INDD-07 | L | 80 | M | FTLD-bvFTD | 1.Argyrophilic grain disease, 2.PSP | 28 |
| INDD-08 | R | 77 | M | Corticobasal syndrome | 1. CBD, 2. FTLD-TDP | 4 |
| INDD-09 | R | 82 | F | Dementia of undetermined etiology | 1.FTLD-TDP, 2.PART | 12 |
| INDD-10 | R | 83 | R | Corticobasal syndrome | 1. PSP, 2. low ADNC | 4 |

**Supplementary Figure 1** Potential landmarks used to anchor the anterior borders of the MTL cortical subregions to.

In each row, in the left image the considered landmark has not yet appeared, the middle image marks the first slide where the landmark appears, and the right one shows the subsequent slide.

The first image of temporal pole (a) marks the first histology slide of the medial temporal lobe. Note that these landmarks are chosen as they are observable on both the histology and MRI.

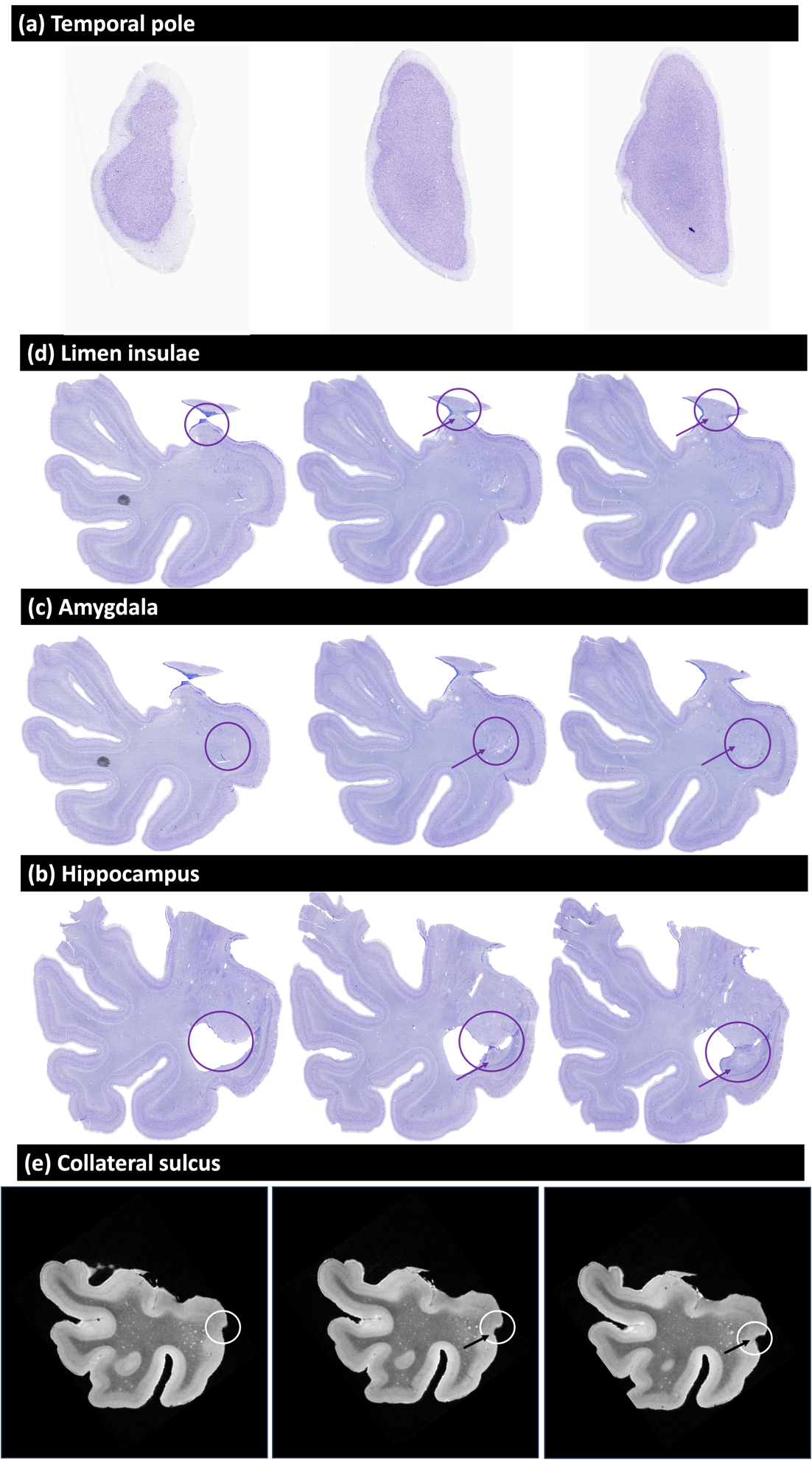
 **Supplementary Figure 2** The measurement of distances on the histology sections using Adobe Illustrator. Black lines are neuroanatomist’s annotations of boundaries between MTL subregions, and the red curve measures the distance between a given boundary (BA35 inferior border) and a given anatomical landmark proposed for consideration in the MRI protocol (fundus of the CS).
Note that histology annotations shown here include fine-grained partitions of BA35 into subregions 35v, 35o, and 35d. These were combined as just BA35 for the development of the protocol.

By measuring the length of the scale bar in Adobe Illustrator units, we were able to convert all the measurements to units of millimeters. In the image, the 5 mm scale bar corresponds to ∼27.5 mm in Adobe Illustrator. Hence the length of the red curve between the CS fundus and inferior border of the BA35 is estimated to be 3.4 mm.

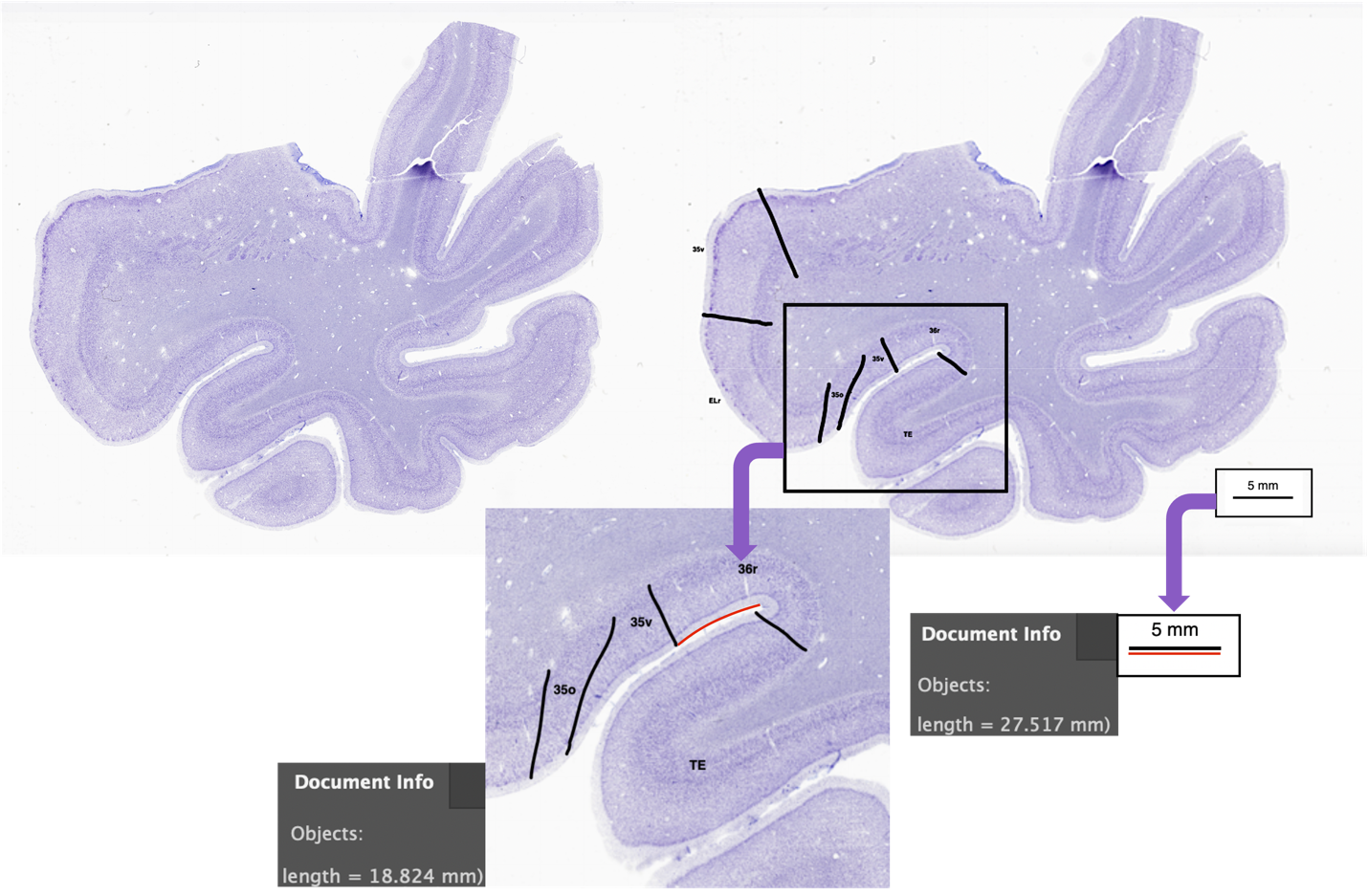

**Supplementary Figure 3** Supplementary Fig 3. Landmarks observable on MRI in coronal slices. Abbreviations: PHG: parahippocampal gyrus; FG: fusiform gyrus; CS: collateral sulcus

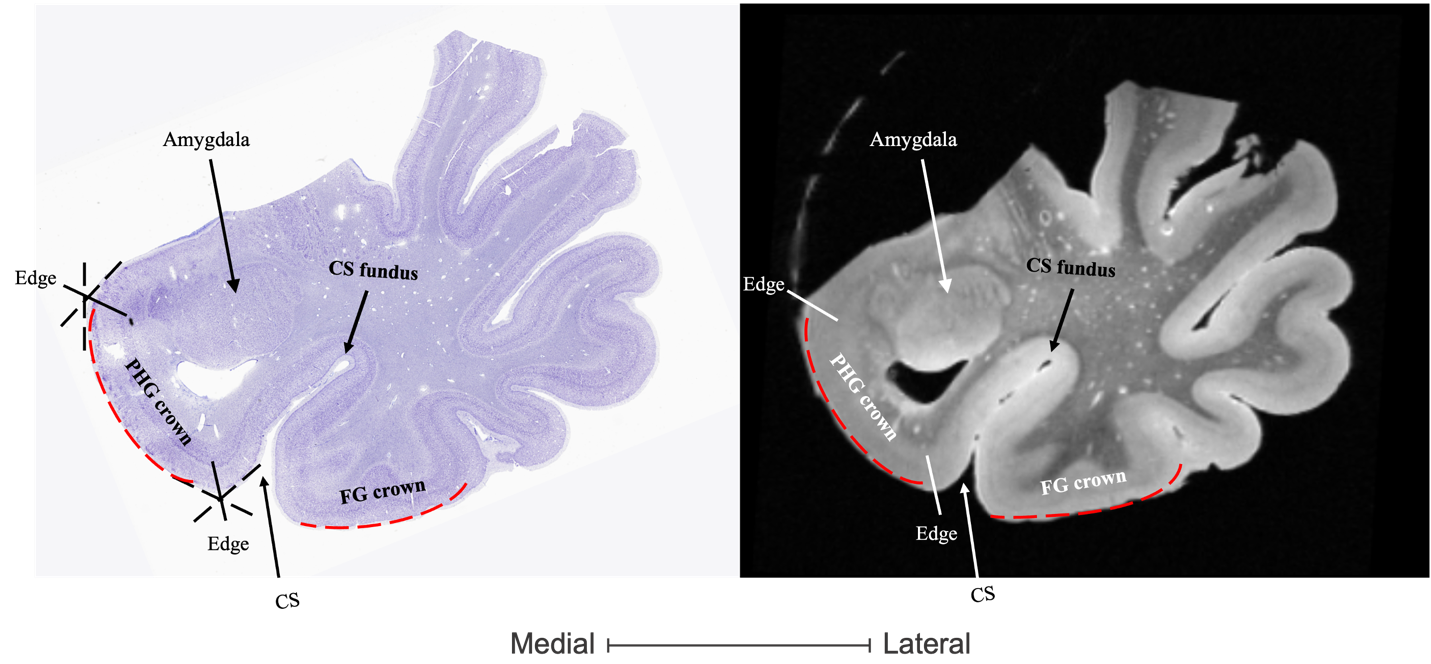

**Supplementary Table 2** Summary of the composition of the in vivo atlas set, including demographics, and cognitive testing. The P‐values are two‐tailed and computed using the t‐test for numerical variables and using the Fisher exact test for sex. Abbreviations: MMSE: Mini‐Mental State Examination; CERAD: Consortium to Establish a Registry for Alzheimer's Disease.

|  | **Atlas subset (*n* = 29)** | | | |  |
| --- | --- | --- | --- | --- | --- |
|  | **NC (*n* = 15)** | | **aMCI (*n* = 14)** | |  |
|  | **Mean ± S.D.** | **Range** | **Mean ± S.D.** | **Range** | ***P*** |
| Sex (male/female) | 7/8 | | 6/8 | | 1.0000 |
| Age | 66.3 ± 9.5 | 54−84 | 71.9 ± 6.2 | 63−80 | 0.0696 |
| Education  (years) | 15.6 ± 2.6 | 12−20 | 16.9 ± 2.8 | 12−20 | 0.1994 |
| MMSE | 29.5 ± 1.0 | 27−30 | 26.9 ± 1.7 | 24−30 | 0.0001 |
| CERAD word list total | 24.7 ± 2.9 | 21−29 | 16.2 ± 3.2 | 11−23 | 0.0000 |
| Delayed recall | 8.7 ± 1.8 | 4−10 | 3.4 ± 2.1 | 0−8 | 0.0000 |

**Supplementary Table 3** Distances from the ERC histological boundaries to landmarks observable on MRI for cases with a deep vs. shallow collateral sulcus and cases with neurodegenerative diseases vs. without neurodegenerative diseases. A negative value reflects the situation where the actual border is located laterally of the landmark and a positive value where the actual border is located medially of the landmark. Abbreviations: CS: collateral sulcus; PHG: parahippocampal gyrus; NDD: neurodegenerative disease

Please note that when comparing cases with deep vs. shallow CS, we only considered borders in the vicinity of the CS to show consistency of the border placement between the two groups in this area.
*The borders outside CS area were marked with N/A.
A cut-off of 7mm in the first slide where the hippocampal head appears was used to determine if cases had a shallow or deep CS(22).

| **ERC** | | | | | | | | | |
| --- | --- | --- | --- | --- | --- | --- | --- | --- | --- |
| **5mm anterior to the hippocampus** | | | | | | | | | |
|  | **Lateral ERC border to** | | | **Medial ERC border to** | | | | | |
|  | **Medial CS edge** | | | **Superior PHG edge** | | | **Halfway point of PHG crown** | | |
| Variable: | Mean | Median | SD | Mean | Median | SD | Mean | Median | SD |
| Deep CS | -0.02 | -1.66 | 4.50 | N/A* | | | | | |
| Shallow CS | -0.09 | -2.03 | 4.10 | N/A* | | | | | |
| With NDD | 0.45 | -1.85 | 4.27 | -2.46 | -0.36 | 5.70 | 7.00 | 6.94 | 6.10 |
| Without NDD | -1.04 | -1.44 | 4.29 | -4.31 | -2.46 | 3.27 | 1.69 | 1.71 | 5.18 |
| **First ERC slide** | | | | | | | | | |
|  | **Lateral ERC border to** | | | **Medial ERC border to** | | | | | |
|  | **Medial CS edge** | | | **Superior PHG edge** | | | **Halfway point of PHG crown** | | |
| Variable: | Mean | Median | SD | Mean | Median | SD | Mean | Median | SD |
| Deep CS | 1.20 | 0.73 | 4.40 | N/A* | | | | | |
| Shallow CS | 0.30 | -1.75 | 4.58 | N/A* | | | | | |
| With NDD | 0.85 | 0.00 | 4.34 | -5.71 | -4.81 | 5.12 | 2.16 | 3.38 | 4.12 |
| Without NDD | 0.63 | -1.03 | 4.72 | -4.30 | -2.60 | 7.65 | -0.05 | -1.04 | 4.20 |
| **4mm anterior to the hippocampus** | | | | | | | | | |
|  | **Lateral ERC border to** | | | **Medial ERC border to** | | | | | |
|  | **Medial CS edge** | | | **Superior PHG edge** | | | **Halfway point of PHG crown** | | |
| Variable: | Mean | Median | SD | Mean | Median | SD | Mean | Median | SD |
| Deep CS | -0.96 | -0.28 | 3.32 | N/A* | | |  |  |  |
| Shallow CS | -0.25 | -0.93 | 3.76 | N/A* | | |  |  |  |
| With NDD | -1.83 | -0.91 | 2.03 | -1.83 | -1.56 | 3.56 |  |  |  |
| Without NDD | 0.73 | 0.00 | 4.27 | -3.45 | -3.19 | 4.52 |  |  |  |
| **2.5mm anterior to the hippocampus** | | | | | | | | | |
|  | **Lateral ERC border to** | | | **Medial ERC border to** | | | | | |
|  | **Medial CS edge** | | | **Superior PHG edge** | | | **Halfway point of PHG crown** | | |
| Variable: | Mean | Median | SD | Mean | Median | SD | Mean | Median | SD |
| Deep CS | -0.83 | -1.12 | 3.22 | N/A* | | |  |  |  |
| Shallow CS | -2.67 | -2.78 | 1.36 | N/A* | | |  |  |  |
| With NDD | -1.87 | -2.11 | 1.26 | -0.28 | -0.94 | 4.16 |  |  |  |
| Without NDD | -1.61 | -2.58 | 3.72 | -0.85 | -1.67 | 2.50 |  |  |  |

**Supplementary Table 4** Distances from BA35 histological boundaries to landmarks observable on MRI for cases with a deep vs. shallow collateral sulcus and cases with neurodegenerative diseases vs. without neurodegenerative diseases. A negative value reflects the situation where the actual border is located laterally of the landmark and a positive value where the actual border is located medially of the landmark.

Abbreviations: CS: collateral sulcus; PHG: parahippocampal gyrus; NDD: neurodegenerative disease

Please note that when comparing cases with deep vs. shallow CS, we only considered borders in the vicinity of the CS to show consistency of the border placement between the two groups in this area.
*The borders beyond CS area were marked with N/A.
A cut-off of 7mm in the first slide where the hippocampal head appears was used to determine if cases had a shallow or deep CS(22).

| **BA35** | | | | | | |
| --- | --- | --- | --- | --- | --- | --- |
| **9mm anterior to the hippocampus** | | | | | | |
|  | **Lateral border of BA35 to** | | | **Medial border of BA35 to** | | |
|  | **CS fundus** | | | **Superior PHG edge** | | |
| variable: | Mean | Median | SD | Mean | Median | SD |
| Deep CS | -1.89 | -0.97 | 4.70 | N/A* | | |
| Shallow CS | -0.77 | 0.00 | 2.81 | N/A* | | |
| With NDD | -0.71 | 0.00 | 5.68 | 9.29 | 8.07 | 5.08 |
| Without NDD | -0.44 | -0.45 | 3.95 | 6.09 | 6.49 | 1.27 |
| **7mm anterior to the hippocampus** | | | | | | |
|  | **Lateral border of BA35 to** | | | **Medial border of BA35 to** | | |
|  | **CS fundus** | | | **Superior PHG edge** | | |
| variable: | Mean | Median | SD | Mean | Median | SD |
| Deep CS | -3.78 | 0.00 | 6.86 | N/A* | | |
| Shallow CS | 0.02 | 0.00 | 1.11 | N/A* | | |
| With NDD | -0.93 | 0.00 | 2.47 | 9.25 | 9.35 | 4.31 |
| Without NDD | -2.72 | 0.00 | 6.95 | 10.48 | 9.96 | 4.18 |
| **5mm anterior to the hippocampus** | | | | | | |
|  | **Lateral border of BA35 to** | | | **Medial border of BA35 to** | | |
|  | **CS fundus** | | | **Superior PHG edge** | | |
| variable: | Mean | Median | SD | Mean | Median | SD |
| Deep CS | -1.11 | 0.00 | 2.77 | N/A* | | |
| Shallow CS | 1.11 | 0.95 | 1.01 | N/A* | | |
| With NDD | 1.12 | 0.60 | 1.24 | N/A* | | |
| Without NDD | -0.87 | 0.00 | 2.64 | N/A* | | |
| **4mm anterior to the hippocampus** | | | | | | |
|  | **Lateral border of BA35 to** | | | **Medial border of BA35** | | |
|  | **CS fundus** | | |  | | |
| Variable: | Mean | Median | SD |  |  |  |
| Deep CS | -0.21 | 0.32 | 2.63 | Lateral border of ERC | | |
| Shallow CS | 1.21 | 1.38 | 1.03 | Lateral border of ERC | | |
| With NDD | 1.31 | 1.12 | 1.32 | Lateral border of ERC | | |
| Without NDD | -0.48 | 0.00 | 2.48 | Lateral border of ERC | | |
| **2.5mm anterior to the hippocampus** | | | | | | |
|  | **Lateral border of BA35 to** | | | **Medial border of BA35** | | |
|  | **CS fundus** | | |  | | |
| Variable: | Mean | Median | SD |  |  |  |
| Deep CS | -0.38 | 0.00 | 2.47 | Lateral border of ERC | | |
| Shallow CS | 1.49 | 1.23 | 1.24 | Lateral border of ERC | | |
| With NDD | 1.27 | 1.16 | 1.01 | Lateral border of ERC | | |
| Without NDD | -0.33 | 0.00 | 2.82 | Lateral border of ERC | | |

**Supplementary Table 5** Distances from BA36 histological boundaries to landmarks observable on MRI for cases with a deep vs. shallow collateral sulcus and cases with neurodegenerative diseases vs. without neurodegenerative diseases. A negative value reflects the situation where the actual border is located laterally of the landmark and a positive value where the actual border is located medially of the landmark.

Abbreviations: CS: collateral sulcus; PHG: parahippocampal gyrus; NDD: neurodegenerative disease

Please note that when comparing cases with deep vs. shallow CS, we only considered borders in the vicinity of the CS to show consistency of the border placement between the two groups in this area.
*The borders beyond CS area were marked with N/A.
A cut-off of 7mm in the first slide where the hippocampal head appears was used to determine if cases had a shallow or deep CS (22).

| **BA36** | | | | | | |
| --- | --- | --- | --- | --- | --- | --- |
| **10mm anterior to the hippocampus** | | | | | | |
|  | **Lateral border of BA36 to** | | | **Medial border of BA36 to** | | |
|  | **CS fundus** | | | **Halfway point of the medial bank of the CS** | | |
| variable: | Mean | Median | SD | Mean | Median | SD |
| Deep CS | 1.83 | 0.00 | 2.59 | 1.03 | 0.57 | 3.92 |
| Shallow CS | 5.06 | 0.53 | 6.93 | -3.71 | 0.23 | 9.89 |
| With NDD | 2.90 | 0.00 | 5.54 | -0.99 | 0.55 | 6.03 |
| Without NDD | 4.13 | 2.36 | 5.36 | -1.77 | 0.57 | 9.83 |
| **2.5 to 9mm anterior to the hippocampus** | | | | | | |
|  | **Lateral border of BA36 to** | | | **Medial border of BA36** | | |
|  | **Halfway point of PHG crown** | | |  | | |
|  | N/A* | | | Lateral border of BA35 | | |

**Supplementary Table 6** Volume comparisons of anterior entorhinal cortex (ERC), Brodmann area 35 (BA35), and Brodmann area 36 (BA36) between cases with deep and shallow collateral sulcus (CS). Volumes were obtained using automatic segmentation, and group differences were assessed using a two-sample t-test. No significant differences were observed across the groups (p > 0.05 for all comparisons).

| Region | Shallow sulcus  Volume (mm^3^)  Mean±SD | Deep sulcus  Volume (mm^3^)  Mean±SD | T  statistic | P  value |
| --- | --- | --- | --- | --- |
| ERC | 717.79±77.68 | 719.90±140.25 | 0.04 | 0.96 |
| BA35 | 997.52±185.49 | 1064.04±133.68 | 1.11 | 0.27 |
| BA36 | 2336.33±550.02 | 2300.57±346.56 | -0.21 | 0.83 |

*Abbreviations: ERC=entorhinal cortex; BA=Brodmann area.*

**Supplementary Figure 4** Modified lateral border of BA36 for the second section of this region located 9 mm anterior to the head of the hippocampus. The green line is the distance from the fundus to the midpoint of the fusiform gyrus. Lateral border of BA36 was placed at the midpoint of this distance to ensure a smoother transformation of borders between different sections.

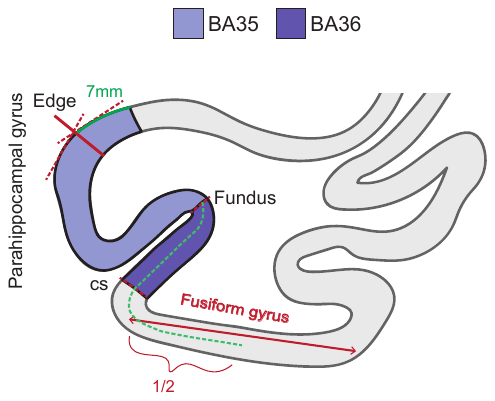

**Supplementary Figure 5** 3D rendering of the anterior regions of the MTL for two cases from our dataset showing the histology annotations mapped into ex vivo MRI space. For both cases the top panel shows all three labels, while the lower two panels focus on the border between either ERC and BA35 or between BA35 and BA36. The figure demonstrates the consistency in border locations between consecutive slices.
Abbreviations: BA=Brodmann area; ERC=entorhinal cortex; HNL: the Human Neuroanatomy Lab; CNDR: the Center for Neurodegenerative Disease Research.

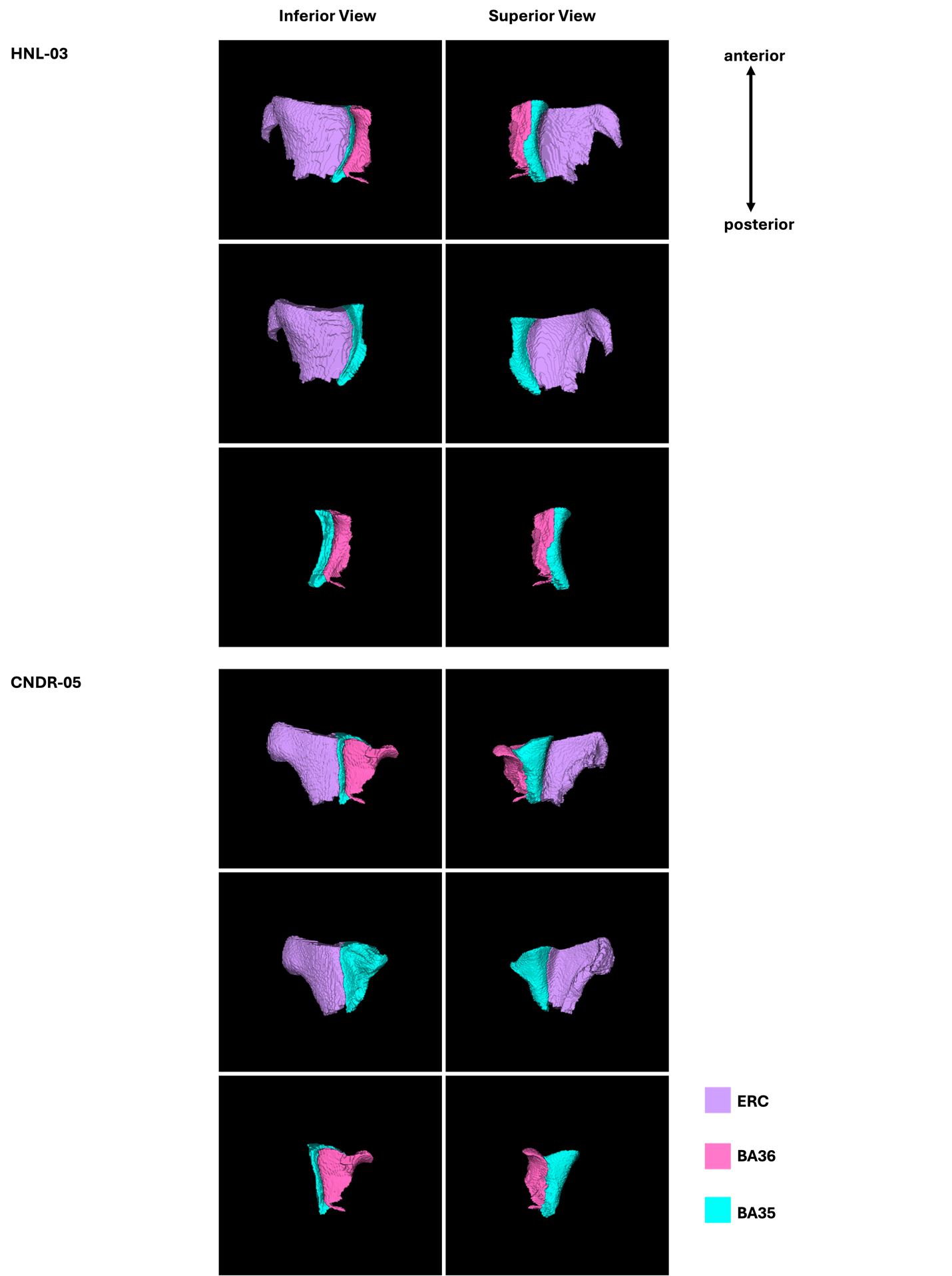

**Supplementary Figure 6** 3D rendering of the segmentation protocol applied manually to two cases from our dataset. For both cases the top panel shows all three labels, while the lower two panels focus on the border between either ERC and BA35 or between BA35 and BA36. The figure demonstrates the consistency in border locations between consecutive slices.
Abbreviations: BA=Brodmann area; ERC=entorhinal cortex; HNL: the Human Neuroanatomy Lab; CNDR: the Center for Neurodegenerative Disease Research; MCI: mild cognitive impairment.

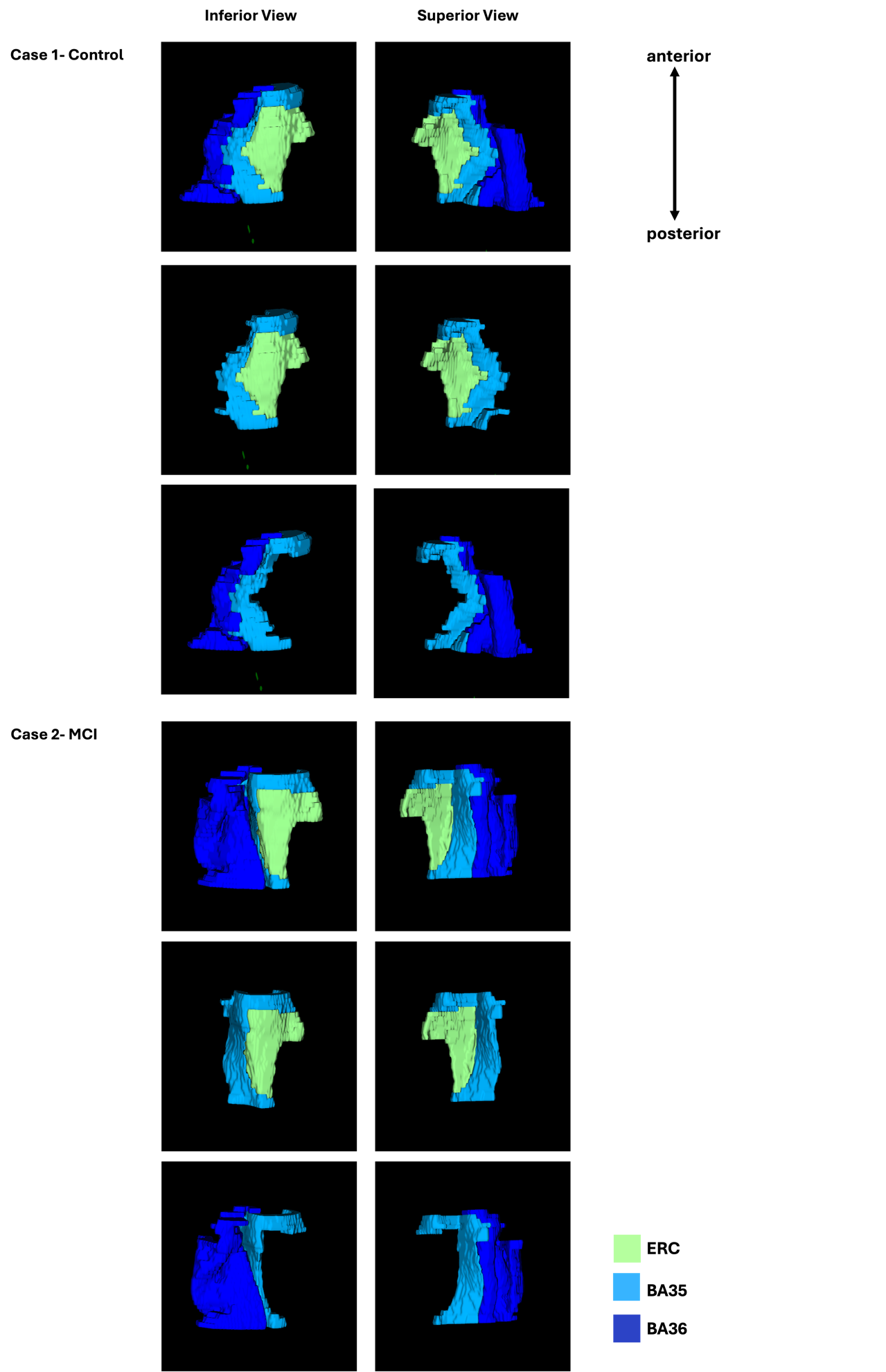

**Supplementary Figure 7** 3D rendering of the automatic segmentation of the anterior regions of the MTL for four cases from the ASHS atlas set, including two control subjects and two individuals with mild cognitive impairment (MCI).

Abbreviations: BA: Brodmann area; ERC: Entorhinal cortex; MCI: mild cognitive impairment.

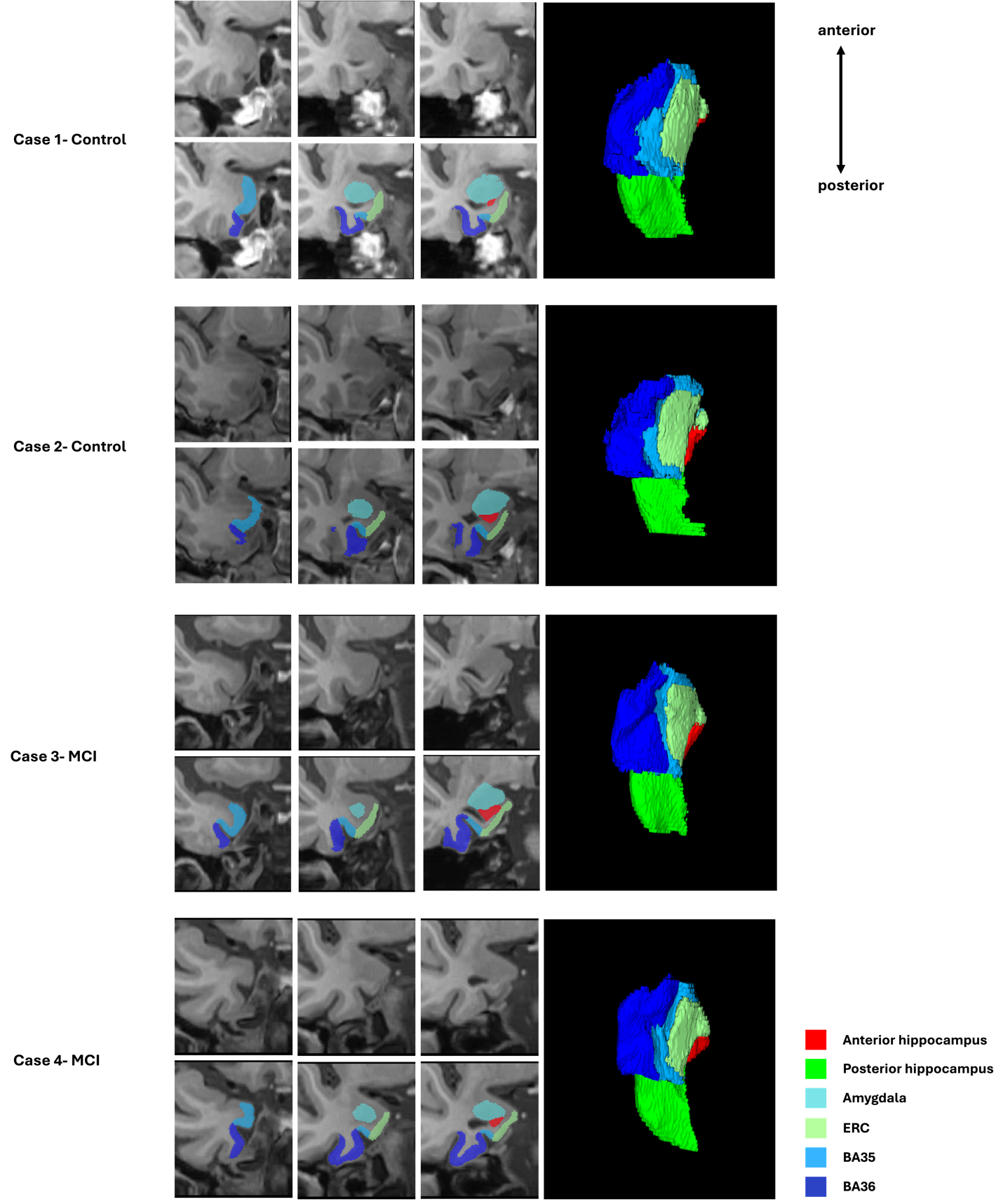
