## Supplementary material for "Developing an anatomically valid segmentation protocol for anterior regions of the medial temporal lobe for neurodegenerative diseases": Supp_fig_1

**Supplementary Figure 1** Potential landmarks used to anchor the anterior borders of the MTL cortical subregions to.

In each row, in the left image the considered landmark has not yet appeared, the middle image marks the first slide where the landmark appears, and the right one shows the subsequent slide.

The first image of temporal pole (a) marks the first histology slide of the medial temporal lobe. Note that these landmarks are chosen as they are observable on both the histology and MRI.

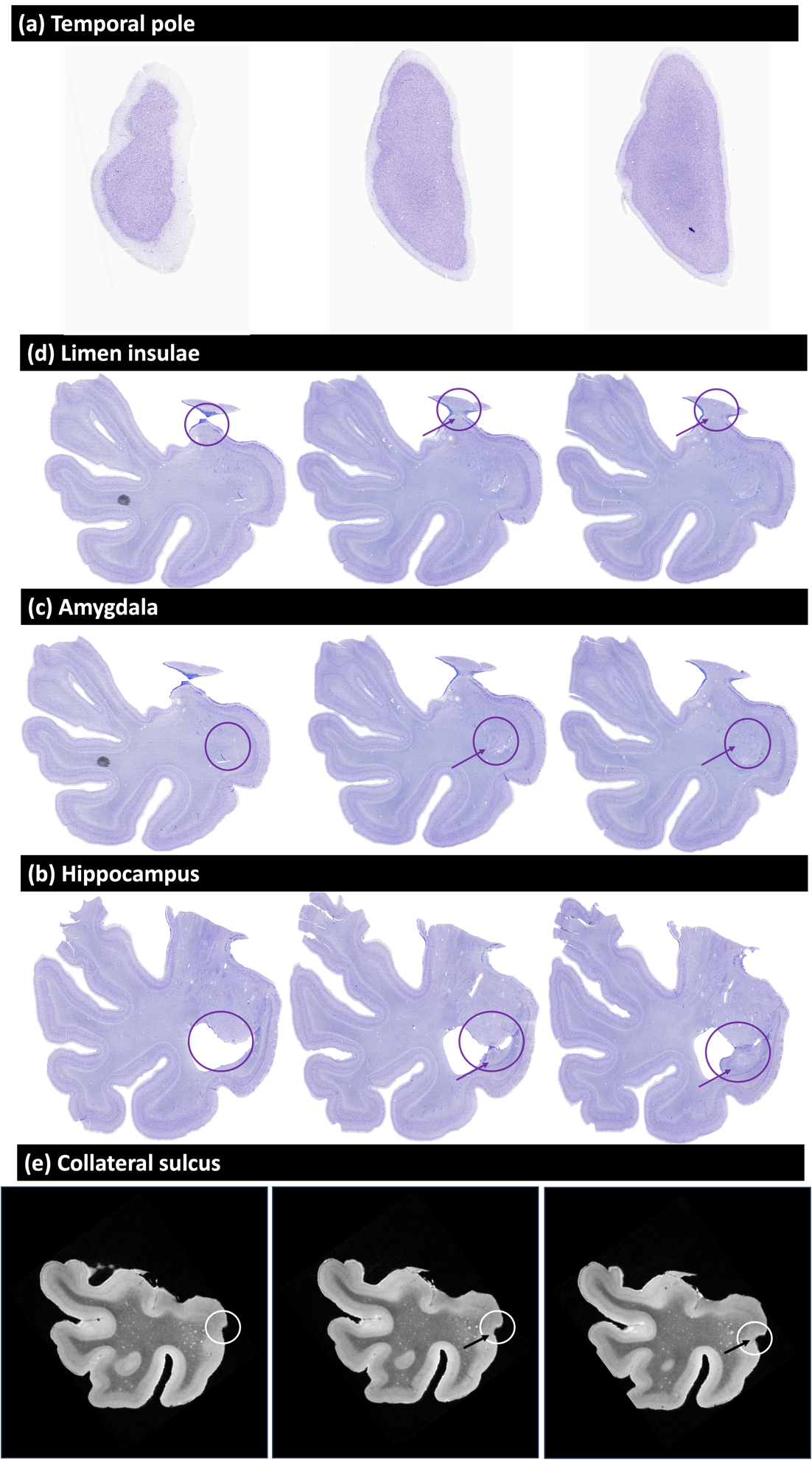
