## Supplementary material for "Developing an anatomically valid segmentation protocol for anterior regions of the medial temporal lobe for neurodegenerative diseases": Supp_fig_2

**Supplementary Figure 2** The measurement of distances on the histology sections using Adobe Illustrator. Black lines are neuroanatomist’s annotations of boundaries between MTL subregions, and the red curve measures the distance between a given boundary (BA35 inferior border) and a given anatomical landmark proposed for consideration in the MRI protocol (fundus of the CS).
Note that histology annotations shown here include fine-grained partitions of BA35 into subregions 35v, 35o, and 35d. These were combined as just BA35 for the development of the protocol.

By measuring the length of the scale bar in Adobe Illustrator units, we were able to convert all the measurements to units of millimeters. In the image, the 5 mm scale bar corresponds to ∼27.5 mm in Adobe Illustrator. Hence the length of the red curve between the CS fundus and inferior border of the BA35 is estimated to be 3.4 mm.

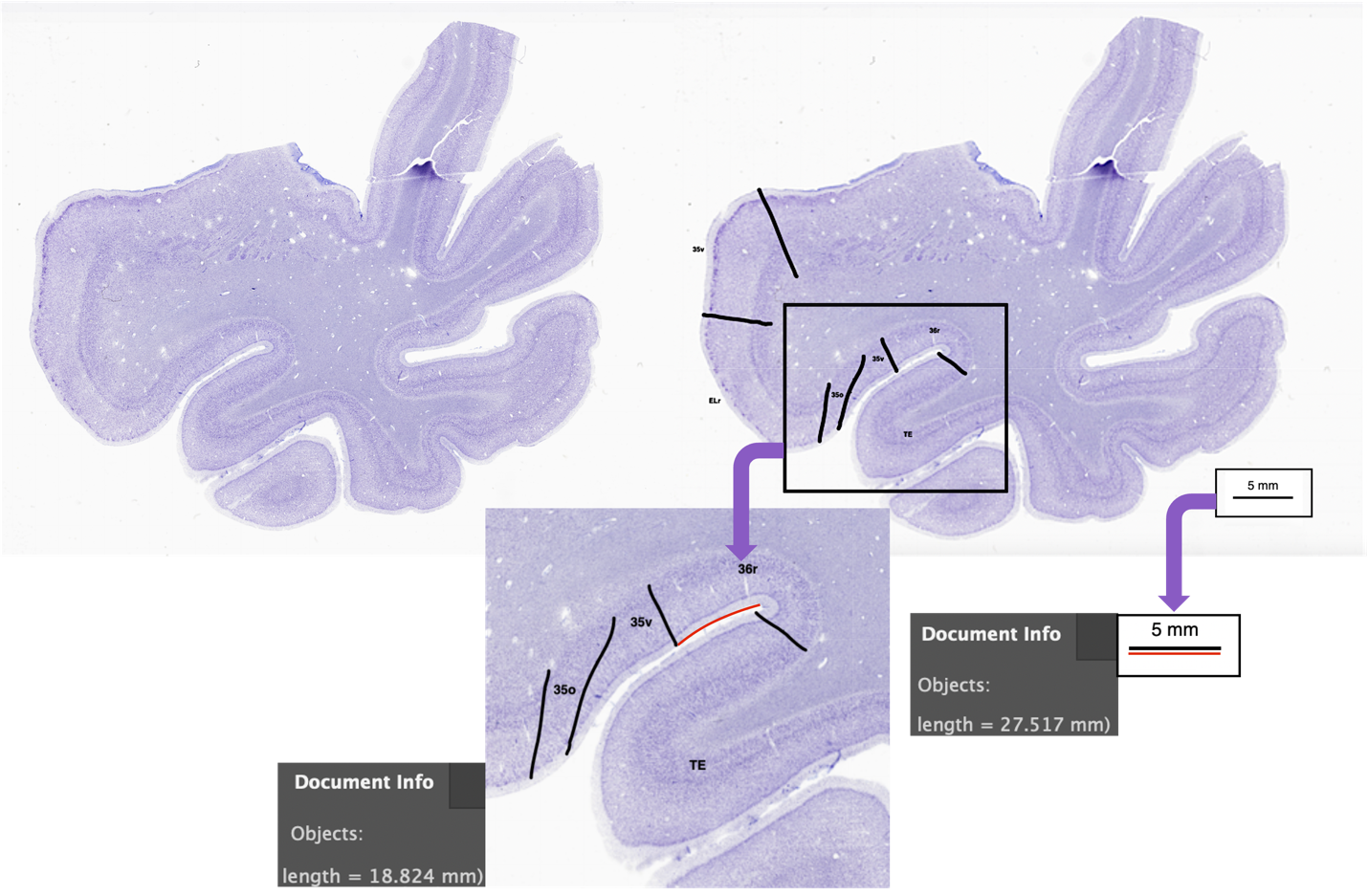
