## Supplementary material for "Developing an anatomically valid segmentation protocol for anterior regions of the medial temporal lobe for neurodegenerative diseases": Supp_fig_3

**Supplementary Figure 3** Supplementary Fig 3. Landmarks observable on MRI in coronal slices. Abbreviations: PHG: parahippocampal gyrus; FG: fusiform gyrus; CS: collateral sulcus

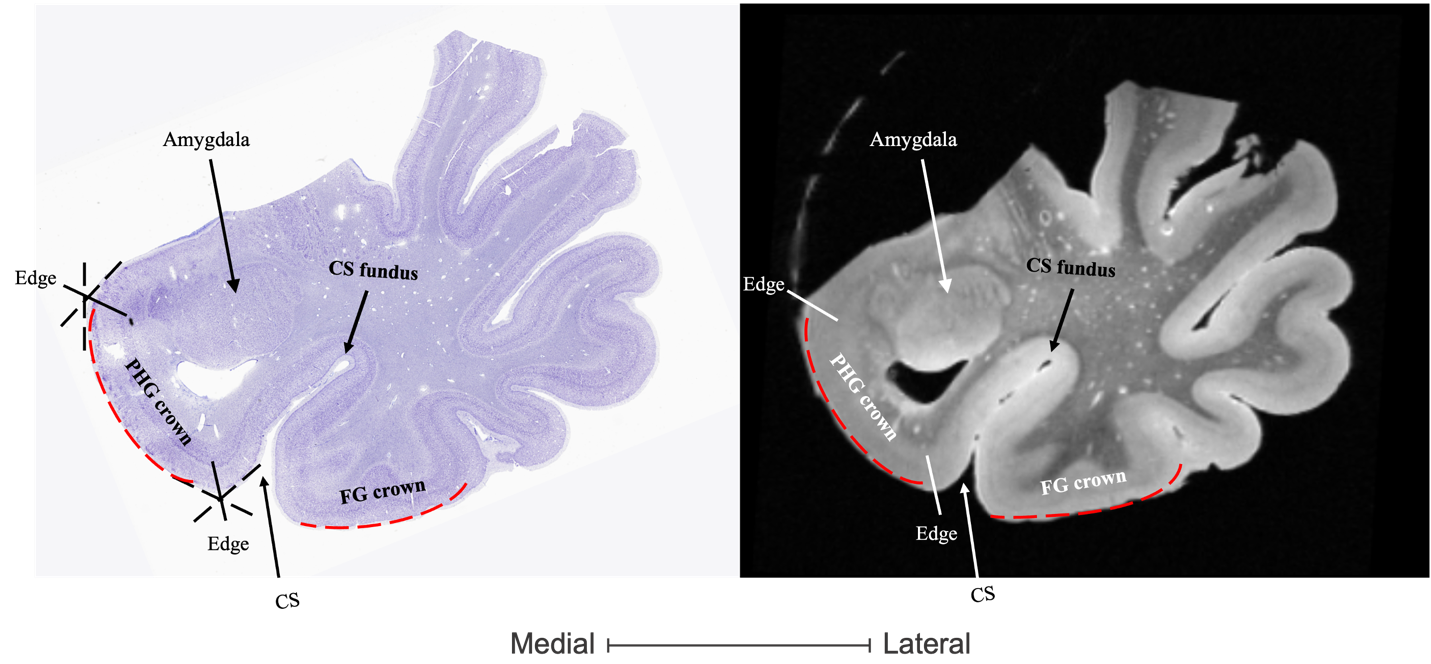
