## Supplementary material for "Developing an anatomically valid segmentation protocol for anterior regions of the medial temporal lobe for neurodegenerative diseases": Supp_fig_4

**Supplementary Figure 4** Modified lateral border of BA36 for the second section of this region located 9 mm anterior to the head of the hippocampus. The green line is the distance from the fundus to the midpoint of the fusiform gyrus. Lateral border of BA36 was placed at the midpoint of this distance to ensure a smoother transformation of borders between different sections.

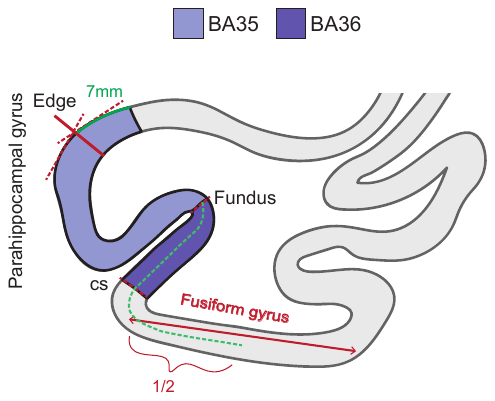
