## Supplementary material for "Developing an anatomically valid segmentation protocol for anterior regions of the medial temporal lobe for neurodegenerative diseases": Supp_fig_6

**Supplementary Figure 6** 3D rendering of the segmentation protocol applied manually to two cases from our dataset. For both cases the top panel shows all three labels, while the lower two panels focus on the border between either ERC and BA35 or between BA35 and BA36. The figure demonstrates the consistency in border locations between consecutive slices.
Abbreviations: BA=Brodmann area; ERC=entorhinal cortex; HNL: the Human Neuroanatomy Lab; CNDR: the Center for Neurodegenerative Disease Research; MCI: mild cognitive impairment.

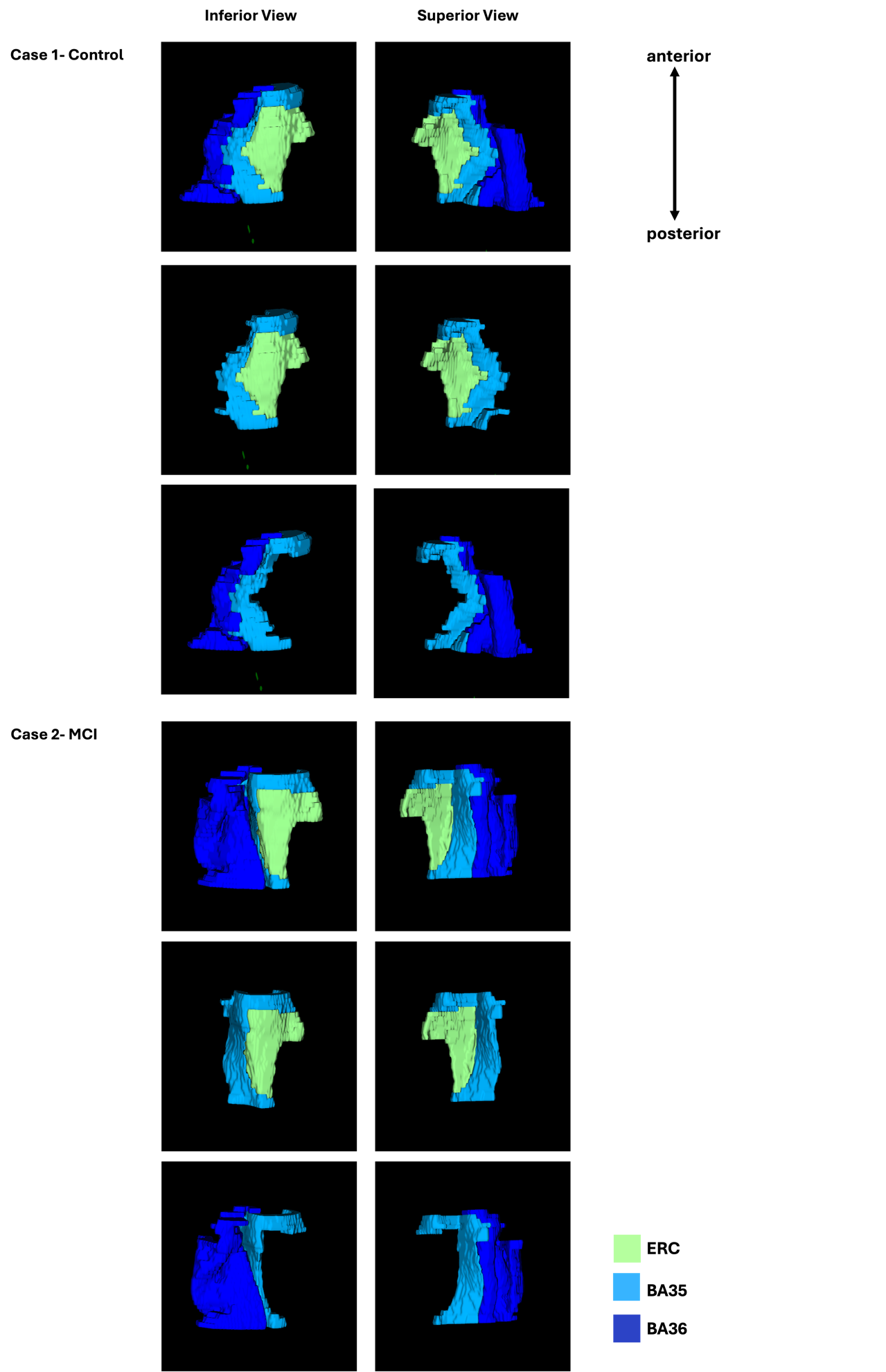
