## Supplementary material for "Developing an anatomically valid segmentation protocol for anterior regions of the medial temporal lobe for neurodegenerative diseases": Supp_fig_7

**Supplementary Figure 7** 3D rendering of the automatic segmentation of the anterior regions of the MTL for four cases from the ASHS atlas set, including two control subjects and two individuals with mild cognitive impairment (MCI).

Abbreviations: BA: Brodmann area; ERC: Entorhinal cortex; MCI: mild cognitive impairment.

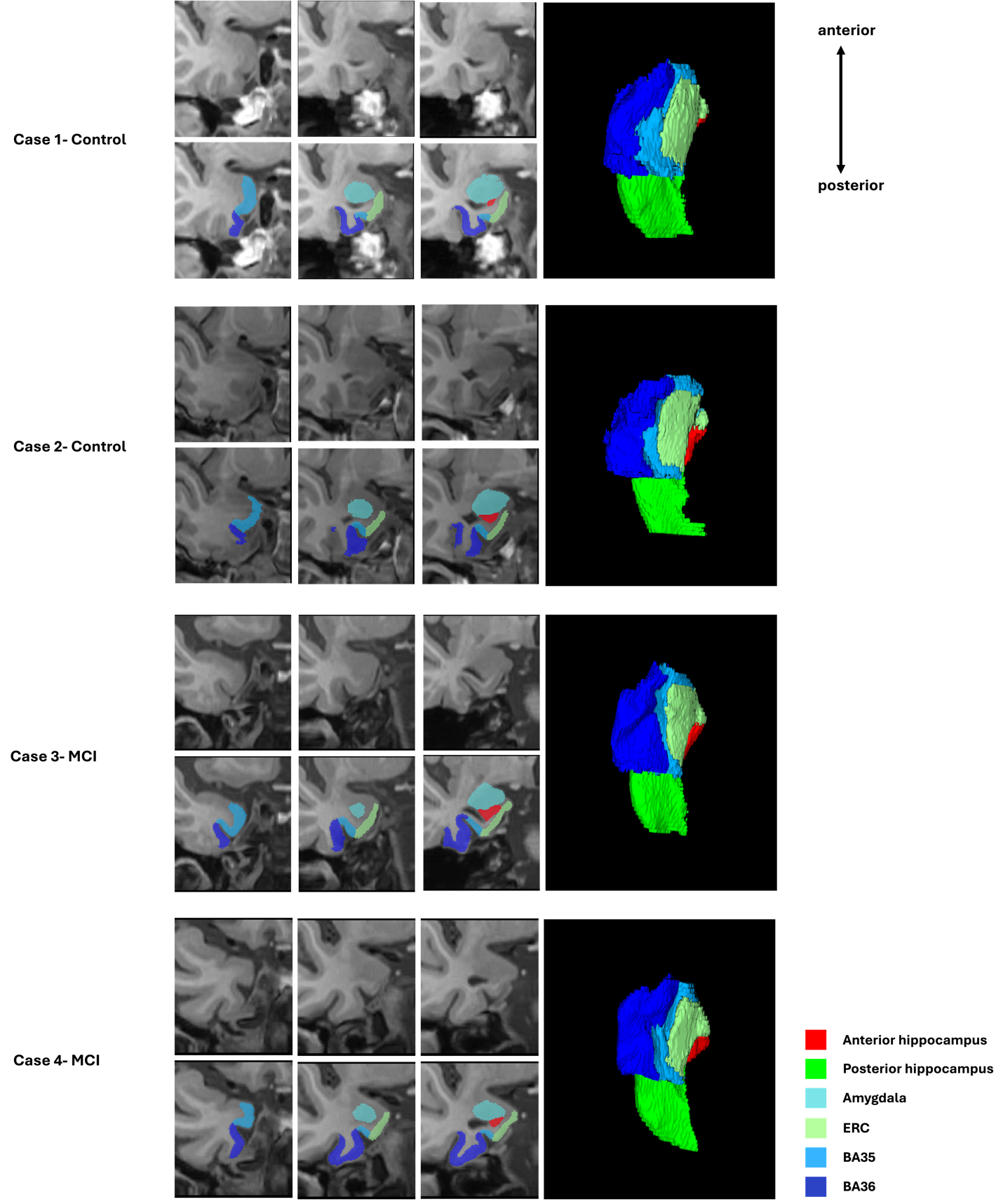
