## Supplementary_tables for "Developing an anatomically valid segmentation protocol for anterior regions of the medial temporal lobe for neurodegenerative diseases"

| **BA35** | | | | | | |
| --- | --- | --- | --- | --- | --- | --- |
| **9mm anterior to the hippocampus** | | | | | | |
|  | **Lateral border of BA35 to** | | | **Medial border of BA35 to** | | |
|  | **CS fundus** | | | **Superior PHG edge** | | |
| variable: | Mean | Median | SD | Mean | Median | SD |
| Deep CS | -1.89 | -0.97 | 4.70 | N/A* | | |
| Shallow CS | -1.09 | 0.00 | 3.02 | N/A* | | |
| With NDD | -2.11 | 0.00 | 4.08 | 9.55 | 8.30 | 5.37 |
| Without NDD | -0.44 | -0.45 | 3.95 | 5.96 | 5.81 | 1.51 |
| **7mm anterior to the hippocampus** | | | | | | |
|  | **Lateral border of BA35 to** | | | **Medial border of BA35 to** | | |
|  | **CS fundus** | | | **Superior PHG edge** | | |
| variable: | Mean | Median | SD | Mean | Median | SD |
| Deep CS | -3.78 | 0.00 | 6.86 | N/A* | | |
| Shallow CS | 0.02 | 0.00 | 1.11 | N/A* | | |
| With NDD | -0.93 | 0.00 | 2.47 | 9.25 | 9.35 | 4.31 |
| Without NDD | -2.72 | 0.00 | 6.95 | 10.48 | 9.96 | 4.18 |
| **5mm anterior to the hippocampus** | | | | | | |
|  | **Lateral border of BA35 to** | | | **Medial border of BA35 to** | | |
|  | **CS fundus** | | | **Superior PHG edge** | | |
| variable: | Mean | Median | SD | Mean | Median | SD |
| Deep CS | -1.11 | 0.00 | 2.77 | N/A* | | |
| Shallow CS | 1.11 | 0.95 | 1.01 | N/A* | | |
| With NDD | 1.12 | 0.60 | 1.24 | N/A* | | |
| Without NDD | -0.87 | 0.00 | 2.64 | N/A* | | |
| **4mm anterior to the hippocampus** | | | | | | |
|  | **Lateral border of BA35 to** | | | **Medial border of BA35** | | |
|  | **CS fundus** | | |  | | |
| Variable: | Mean | Median | SD |  |  |  |
| Deep CS | -0.21 | 0.32 | 2.63 | Lateral border of ERC | | |
| Shallow CS | 1.21 | 1.38 | 1.03 | Lateral border of ERC | | |
| With NDD | 1.31 | 1.12 | 1.32 | Lateral border of ERC | | |
| Without NDD | -0.48 | 0.00 | 2.48 | Lateral border of ERC | | |
| **2.5mm anterior to the hippocampus** | | | | | | |
|  | **Lateral border of BA35 to** | | | **Medial border of BA35** | | |
|  | **CS fundus** | | |  | | |
| Variable: | Mean | Median | SD |  |  |  |
| Deep CS | -0.38 | 0.00 | 2.47 | Lateral border of ERC | | |
| Shallow CS | 1.49 | 1.23 | 1.24 | Lateral border of ERC | | |
| With NDD | 1.27 | 1.16 | 1.01 | Lateral border of ERC | | |
| Without NDD | -0.33 | 0.00 | 2.82 | Lateral border of ERC | | |

**Supplementary Table 5** Distances from BA36 histological boundaries to landmarks observable on MRI for cases with a deep vs. shallow collateral sulcus and cases with neurodegenerative diseases vs. without neurodegenerative diseases. A negative value reflects the situation where the actual border is located laterally of the landmark and a positive value where the actual border is located medially of the landmark.
